# A cholinergic mechanism orchestrating task-dependent computation across the cortex

**DOI:** 10.1101/2025.11.26.690825

**Authors:** Jiaqi Keith Luo, Junhua Tan, Erin Myhre, Peter S. Salvino, Amy P. Rapp, Lucas Pinto

## Abstract

In an ever-changing environment, animals often need to switch between performing different tasks involving distinct sets of cognitive processes. Many such tasks involve neural activity distributed across the cortex, with dynamics that depend on both task demands and behavioral strategy^1–17^. A fundamental but unanswered question is what circuit mechanisms orchestrate these task-dependent dynamics. Here, we hypothesized that acetylcholine release in the cortex plays a key role. The cortex’s only long-range source of this neuromodulator is the basal forebrain cholinergic system, which targets the entire cortical sheet and can individually modulate single regions on sub-second timescales^18–26^. To test our hypothesis, we first imaged cholinergic axons innervating the cortex while mice switched frequently between two navigational decision-making tasks in virtual reality (VR), only one of which required gradual accumulation of sensory evidence. We found that cholinergic input to the cortex is spatiotemporally heterogeneous and multiplexes sensory, motor, arousal, and cognitive signals in a task- and strategy-dependent fashion, with overall higher activity during evidence accumulation. Crucially, beyond contextual variables, cholinergic activity directly tracked task computations themselves, encoding an evidence-dependent decision variable only in the accumulation task. To test if acetylcholine release is causal to the performance of each task, we optogenetically silenced cholinergic terminals in the cortex while simultaneously imaging excitatory cortical activity. We found that this input is selectively required for evidence accumulation, and for large-scale cortical coding of evidence and choice during the accumulation task. Thus, we have identified a new cholinergic mechanism that orchestrates cortex-wide activity in a task-dependent manner and serves as a key node in the distributed brain network underlying the accumulation of sensory evidence.

## Introduction

Basal forebrain cholinergic neurons and their cortical projections have been implicated in multiple aspects of behavior, including: arousal^25,27–33^; sensory processing, attention and perceptual decision making^21–23,26,31,33–43^; action^24,25,38,44–47^; and reinforcement, learning and plasticity^22,24,38,47– 52^. However, many cognitive tasks involve all the different aspects of behavior listed above, which begs the question of whether and how the cholinergic system integrates these signals task dependently. This is likely enabled by the multiplexing of these signals on multiple timescales and by the fact that cholinergic projections to the cortex have a rough topographical arrangement^53–59^ that allows it to modulate cortical dynamics with regional specificity^22–25,31,41,59^. This results in spatiotemporally heterogeneous patterns of cholinergic release across the cortex, at least in the context of modality-specific sensory processing and simple behaviors^21,24,25,30,60^. The consequences of this arrangement for flexible cognitive behavior remain poorly understood.

Here, we hypothesized that task-specific, spatiotemporally heterogeneous cholinergic modulation is a key mechanism that rapidly reorganizes cortex-wide activity to allow it to support different cognitive behaviors, even within a single sensory modality. We took advantage of two tasks for mice navigating in virtual reality (VR), which are similar in their sensorimotor components — known major drivers of cholinergic activity — but differ in their underlying cognitive processes^5,61^. Critically, these two tasks engage different patterns of large-scale cortical dynamics^5^. Specifically, when stimuli must be gradually accrued towards a decision, more distributed cortical activity is required for task performance and the dynamics of different regions are less correlated. Although acetylcholine has been broadly implicated in sensory processing, we hypothesized that it would be especially important for evidence accumulation because it decorrelates cortical activity on microcircuit and cortex-wide scales^23,36,62–65^. To test this, we adapted the two tasks above into a paradigm in which mice switched rapidly between them, allowing us to probe the coding properties and causality of task-specific cholinergic activity across the cortex.

## Results

### A task-switching paradigm for mice navigating in virtual reality

We trained mice to switch dozens of unpredictable times between an evidence-accumulation task and a simpler visual discrimination task happening within the same virtual T-maze, dubbed ‘towers’ and ‘visually guided’ tasks, respectively (**Fig. 1a,b**, 31 ± 6 task blocks per session, 10 ± 2 trials per block for the towers task, 6 ± 1 trials for the visually guided task; mean ± s.d.). To isolate potential neural and behavioral signals related to task context, both tasks started with a context region in which wallpaper gratings of different orientations indicated task identity (50 cm). In the towers task, mice then entered an evidence region (150 cm) in which salient white stimuli (towers) flashed briefly along maze walls (200 ms), with random counts and positions across trials. After a delay region (100 cm), mice were rewarded for turning into the arm on the side with the highest count of tower stimuli. In the visually guided task, mice still experienced the briefly flashing tower stimuli, but throughout the maze the mice could see a tall visual guide indicating the rewarded side arm (250 cm, corresponding to evidence and delay regions in the towers task). Crucially, in many trials the visual guide lay on the side opposite the majority of tower stimuli, allowing for an unambiguous read-out of the mice’s strategy.

**Fig. 1.**
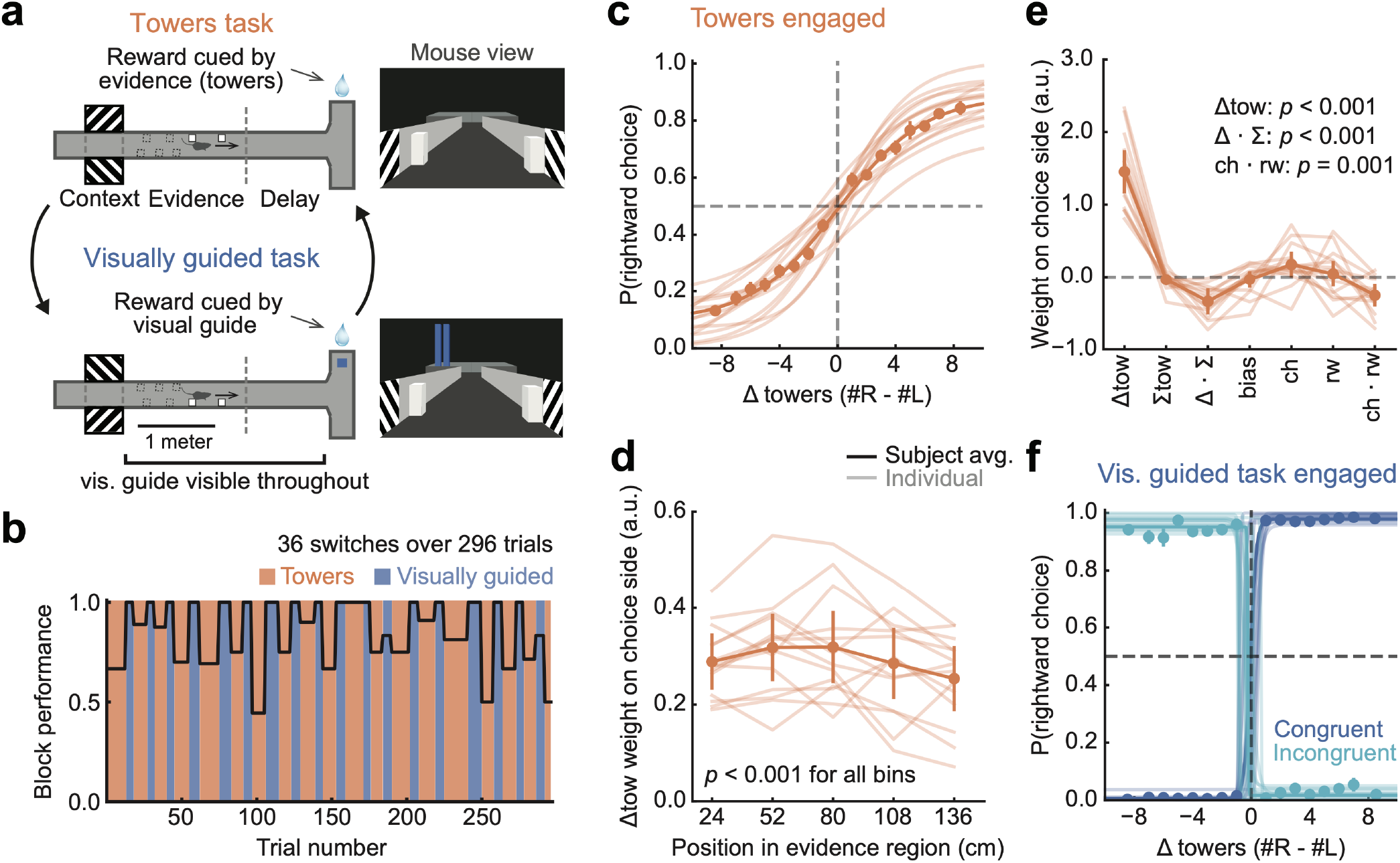
A task-switching paradigm for mice navigating in virtual reality. **a**, Schematics of the maze and the two tasks. **b**, Single-session example of overall task performance. Vertical colored bands: task blocks, black line: block-wise performance. **c**, Psychometric functions showing the fraction of rightward choices as a function of the strength of right-side sensory evidence (Δ towers, #R - #L), for engaged-strategy trials in the towers task. Thin lines: best-fitting sigmoids for individual mice (n = 14), circles: average performance across mice (n = 14 mice, 18,969 trials), thick line: best-fitting sigmoid for average data over subjects, error bars: s.e.m. across mice. **d**, Coefficients from a mixed-effects logistic-regression model predicting choice side from net sensory evidence (Δ towers) from different portions of the evidence region, indicating that the mice integrate evidence from across the maze. The model was fitted using the same trials as panel c. Thin lines: random-effect estimates for individual mice, thick lines: population fixed-effect estimates, error bars: mean ± c.i.. **e**, Coefficients from a similar logistic-regression model as in d, but using different evidence and trial-history terms. Conventions as in d. Δtow: Δ towers, ∑tow: total number of towers (#R + #L), Δ · ∑: interaction between Δtow and ∑tow, bias: side bias, ch: previous choice side, rw: previous reward, ch · rw: interaction between ch and rw, i.e., ‘win-stay/lose-switch’. Printed *p*-values are from a mixed-effects logistic regression. **f**, Psychometric functions for engaged trials in the visually guided task (n = 14 mice, 21,771 trials), plotted separately for congruent and incongruent trials (17,397 and 4,374 trials, respectively) — i.e., ones with the visual guide on the same or opposite side to the one with majority towers, respectively. Conventions as in c.

The mice successfully performed both tasks (n = 14 mice, 286 sessions, 69,677 trials). We used a framework combining generalized linear models and hidden Markov models^66–68^ (GLM-HMM) to separate blocks of trials for each task in which the mice adopted the expected stimulus-driven behavioral strategies (‘engaged’) from degenerate strategies that relied heavily on trial history and side bias in the towers task, or tower-stimulus counts during the visually guided task (‘disengaged’) (**Fig. S1**). During engaged strategies, towers-task performance scaled with the strength of sensory evidence (#Right - #Left towers, or Δ towers) (**Fig. 1c**, 18,969 trials), corresponding to even integration of stimuli from across the evidence region, with little influence of trial history (**Fig. 1d,e**). Conversely, engaged performance of the visually guided task depended only on the side of the visual guide, with mice successfully ignoring the towers during incongruent trials, even when the tower stimuli strongly indicated the opposite choice (**Fig. 1f**, 17,397 congruent and 4,374 incongruent trials respectively). Thus, unless otherwise stated, all subsequent analyses are performed on engaged-strategy trials.

Because the cholinergic system has been implicated in arousal and running-related modulation of cortical activity^44,45,69,70^, we also measured pupil area to estimate arousal levels (n = 11 mice had pupil tracking, 13,479 and 12,999 engaged trials from the towers and visually guided task, respectively), as well as running velocity. We found largely similar pupil dynamics and running patterns across tasks during the evidence region of the maze, when accumulation occurs task dependently (**Fig. S2a,b**, note that the luminance profiles of the tasks are very similar by design). Interestingly, however, we did see small-magnitude differences in speed and pupil diameter between tasks in other maze regions. For example, mice ran slightly but consistently slower in the context region of the towers task, suggesting that they understood and distinguished the task-identity cue (**Fig. S2a**). Moreover, pupil dynamics significantly differed in the first trial of a task block, resembling more the task in the previous block (**Fig. S2c,d**). This could be analogous to the well-documented phenomenon of task-set inertia in humans^71^.

### Cholinergic input to the cortex is strategy and task dependent

We first asked how cholinergic input varies across tasks and cortical regions. We transgenically expressed the Ca^2+^ indicator jGCaMP8m in cholinergic neurons (ChAT-IRES-Cre x flex-jGCaMP8m cross, n = 8 mice, **Figs. 2a, S3a**, with negligible expression of jGCaMP8m in choline acetyltransferase-positive [ChAT+] interneurons in the cortex, **Fig. S3b**). We then used mesoscale widefield Ca^2+^ imaging through the intact cleared skull to measure activity of cholinergic axons from the basal forebrain simultaneously across many areas of the dorsal cortex while mice engaged in task switching (**Fig. 2b**, ∼50µm/pixel, n = 8 mice, 10,099 and 10,945 engaged-strategy trials in the towers and visually guided tasks, respectively).

**Fig. 2.**
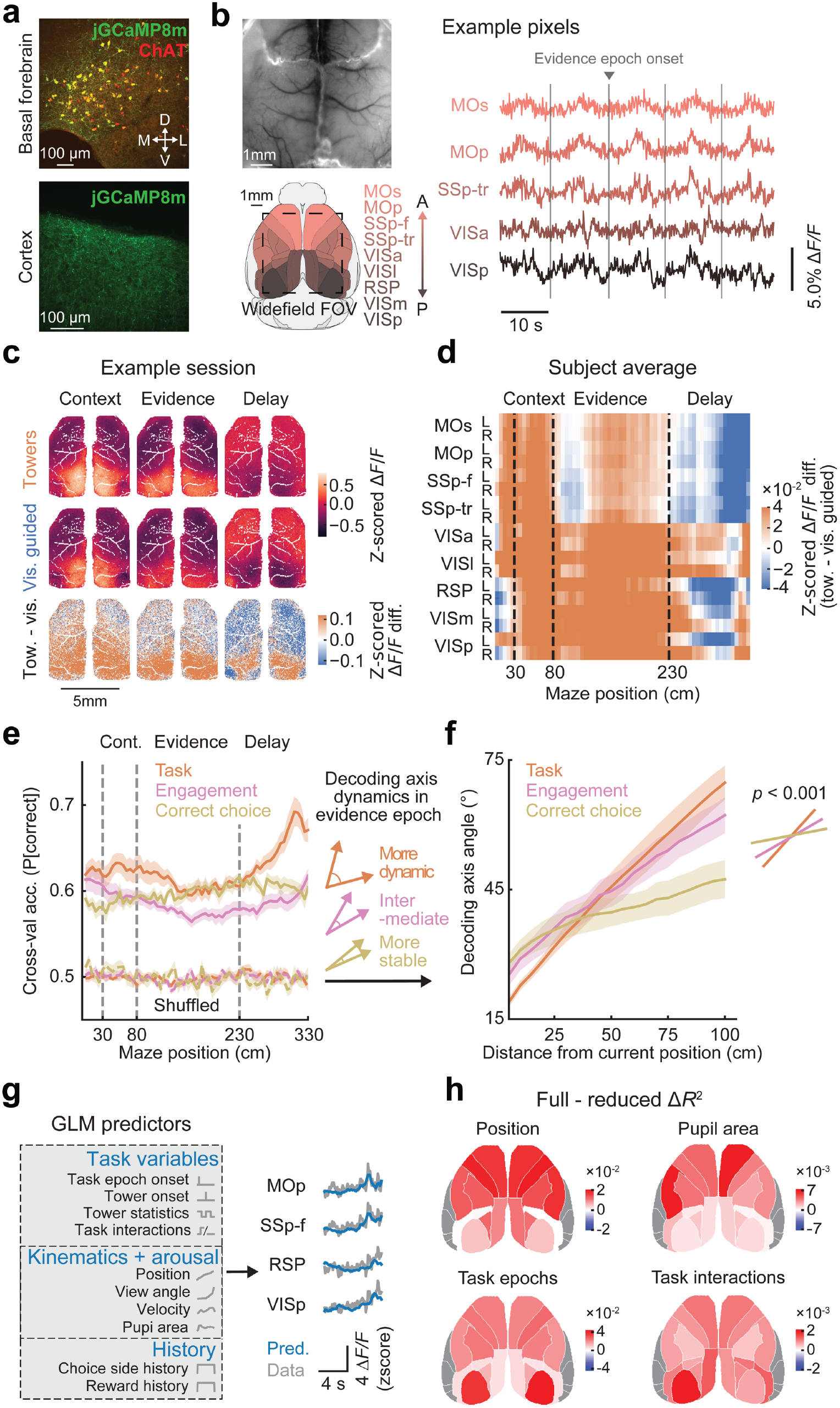
Task- and strategy-specific cholinergic input to the cortex. **a**, Top: Example immunostaining illustrating high overlap between jGCaMP8m (green) and ChAT+ neurons in the basal forebrain (red). Bottom: cholinergic axons innervating the cortex, expressing jGCaMP8m. **b**, Top left: example cleared skull with fluorescence from cholinergic axons. Bottom left: schematics of simultaneously imaged cortical regions, parsed according to the Allen Brain Atlas (ccf3.0). Outline indicates approximate field of view (FOV) for widefield recordings. Right: example Δ*F/F* traces from cholinergic axons. **c**, Top and middle: cholinergic-axon activity maps for an example session, plotted separately for engaged trials in the towers and visually guided tasks (top and middle), and averaged within the different maze regions. Bottom: difference in average activity between the tasks. **d**, Grand average of activity difference between tasks across maze positions (columns), for cholinergic axons innervating different anatomically defined cortical regions (rows) (n = 8 mice, 21,044 trials). **e**, Average cross-validated accuracy of decoders trained to classify the current task, engaged vs. disengaged strategy in the towers task, or whether the upcoming choice was correct in the towers task, from area-averaged cholinergic-axon activity at different maze positions. All trials were used for strategy decoding, and only engaged-strategy trials of the towers task for task and outcome. Shaded regions: ± s.e.m. (n = 8 mice). All positions have significantly higher decoding accuracy than shuffled data (*p* < 0.001 for all positions, t-test with significant *p*-values for at least 3 consecutive positions). **f**, Average decoding-axis angles over position within the evidence region, relative to starting position, and computed within decoders of task, strategy and correct choice. Steeper slopes indicate more dynamic (shorter-timescale) codes. Shaded regions: ± s.e.m. (n = 8 mice). Printed *p*-value is from one-way repeated measure ANOVA on linear fits of slopes and post hoc pairwise tests, with false discovery rate (FDR) correction. **g**, Left: schematics of the linear encoding model of cholinergic activity. Right: examples of single-trial traces in the held-out test set, with model predictions. **h**, Example maps of contributions of different classes of model predictors to explaining cholinergic activity, estimated as the change in the amount of cross-validated variance explained between the full model and a reduced model lacking that set of predictors and refitted to the data.

We observed spatiotemporally dynamic patterns of cholinergic-axon activity during engaged strategies in each task. For instance, in both tasks, cholinergic activity was highest in retrosplenial and posterior parietal areas in the task-context region, in visual and retrosplenial regions during evidence presentation, and in frontal and somatosensory regions as the animals approached the side arms (**Figs. 2c, S4a**). Interestingly, however, when we subtracted visually guided from towers-task cholinergic activity, we found significant differences across cortical regions and maze positions (**Figs. 2c,d, S4b**). Activity across the cortex was higher during the context and evidence regions of the towers task, the latter especially in posterior cortical regions. Conversely, compared to the visually guided task, cholinergic-axon activity across the cortex was lower in the delay region of the towers task. This could reflect stronger reward-expectation signals in the visually guided task^48^, or a short-term-memory-related reduction of cholinergic input in the towers task. To further quantify if task and other behavioral-context information is encoded by cholinergic-activity patterns, we trained logistic-regression decoders from cortex-wide cholinergic-axon activity in different parts of the maze. We used this approach to classify which task the animal was performing, whether the animal was in the engaged strategy, and whether its upcoming choice was correct (the latter two during the towers task). This revealed that cholinergic activity multiplexes all three variables across maze positions (**Fig. 2e**). This was also true when we matched the data by previous-trial outcome to control for potential confounds introduced by different levels of overall performance of each task (**Fig. S4c**). To ask how this encoding of task information changed across the maze, we computed the pairwise angles between the decoding axes for different positions (i.e., the vectors of decoding weights). We then compared the rate at which information changed for task, engagement, and correct choice. Interestingly, the decoding angles diverged most rapidly for task, indicating a highly dynamic code, and slowest for correct choice (**Fig. 2f, S4d**). This indicates that cholinergic input to the cortex multiplexes contextual cognitive variables on multiple timescales, in addition to previously known multiplexing of sensory and motor variables^25,26,31^.

Previous literature suggests that cholinergic-axon activity in the cortex should also track arousal levels, as well as sensory and motor events. To jointly estimate the relative contributions of different behavioral features to cholinergic activity, we fitted L2-regularized linear encoding models to activity averaged within anatomically defined regions, with cross-validation (**Fig. 2g**). Our model predicted held-out data with high accuracy (**Fig. S5a**). Inspection of the fitted coefficients and response kernels confirmed that cholinergic signals indeed reflected multiple cognitive and non-cognitive variables, compatible with previous reports^31,52^ (**Fig. S5b,c**). Further, the pattern of encoding depended on cortical area. For example, cholinergic responses to the onset of tower stimuli were much larger in posterior than frontal regions, compatible with previous work^21,24^ (**Fig. S5c**). To quantify the relative contributions of different task variables to the activity of cholinergic axons innervating different cortical regions, we removed sets of predictors from the model and refitted them, computing the change in variance explained between the full and these reduced models (**Figs. 2h, S5d; Table S1**). This confirmed that different cortical regions received heterogeneous mixtures of task-related signals from cholinergic axons. For example, activity was more impacted by the mouse’s position in the maze and pupil area (an index of arousal) in frontal regions. On the other hand, task interactions were present across the cortex, reflecting the fact that the encoding of task variables was different between the two tasks. Altogether, these results extend previous findings of multiplexed sensory, motor and arousal cholinergic signals to show for the first time that they also reflect cognitive differences between decision-making tasks and behavioral strategies within a task, with higher cholinergic activity when mice gradually accumulate sensory evidence.

### Cholinergic input to the cortex encodes choice formation

Our results so far show that cholinergic input to the cortex dynamically integrates task variables to provide highly contextual signals to different cortical regions. Previous theoretical accounts of neuromodulation had in fact proposed that they act as a permissive input to context-dependent cortical computation^72,73^. However, far beyond just contextual signaling, next we show that cholinergic input reflects the ongoing computation itself. We first used logistic-regression decoders to classify the upcoming choice of the animals from area-level cholinergic activity at different maze positions. To dissociate choice from evidence, we focused on the comparison between the towers task and the incongruent trials of the visually guided task. In the towers task, choice and evidence are decorrelated by the presence of error trials and, in the visually guided task, we used the trials in which the visual pulses (towers) are incongruent with the correct choice. We could decode choice with high accuracy in both tasks, regardless of behavioral strategy, with monotonically increasing levels during and after evidence presentation (**Fig. 3a**). However, engaged strategies in the two tasks involved distinct choice-coding schemes — the angles between choice-decoding axes were orthogonal in the evidence region of the maze (**Figs. 3b, S6a**). We hypothesized that this was in part because choice signals depended on sensory evidence specifically in the towers task. To test this, we projected cortex-wide cholinergic activity during that task onto the decoding axes, separately for each trial. This has been previously interpreted as a moment-by-moment read-out of the amount of choice information, i.e., the decision variable in the task^74–78^. We then grouped trials based on evidence strength (|Δ towers|) and the side with most tower stimuli. Strikingly, during engaged strategies in the towers task, choice information on average gradually ramped up proportionally to evidence strength (**Fig. 3c**). To quantify this, we computed the area under the curve (AUC) for each amount of sensory evidence (**Fig. 3d**). During accumulation, choice information was linearly proportional to evidence strength. Moreover, this effect was specific to both task and within-task strategy: choice signals were insensitive to evidence strength in disengaged towers-task strategies, and only tracked choice side in incongruent visually guided trials (**Fig. 3d**; these effects were true even when regressing out contributions of virtual view angle to cholinergic activity, **Fig. S6b**).

**Fig. 3.**
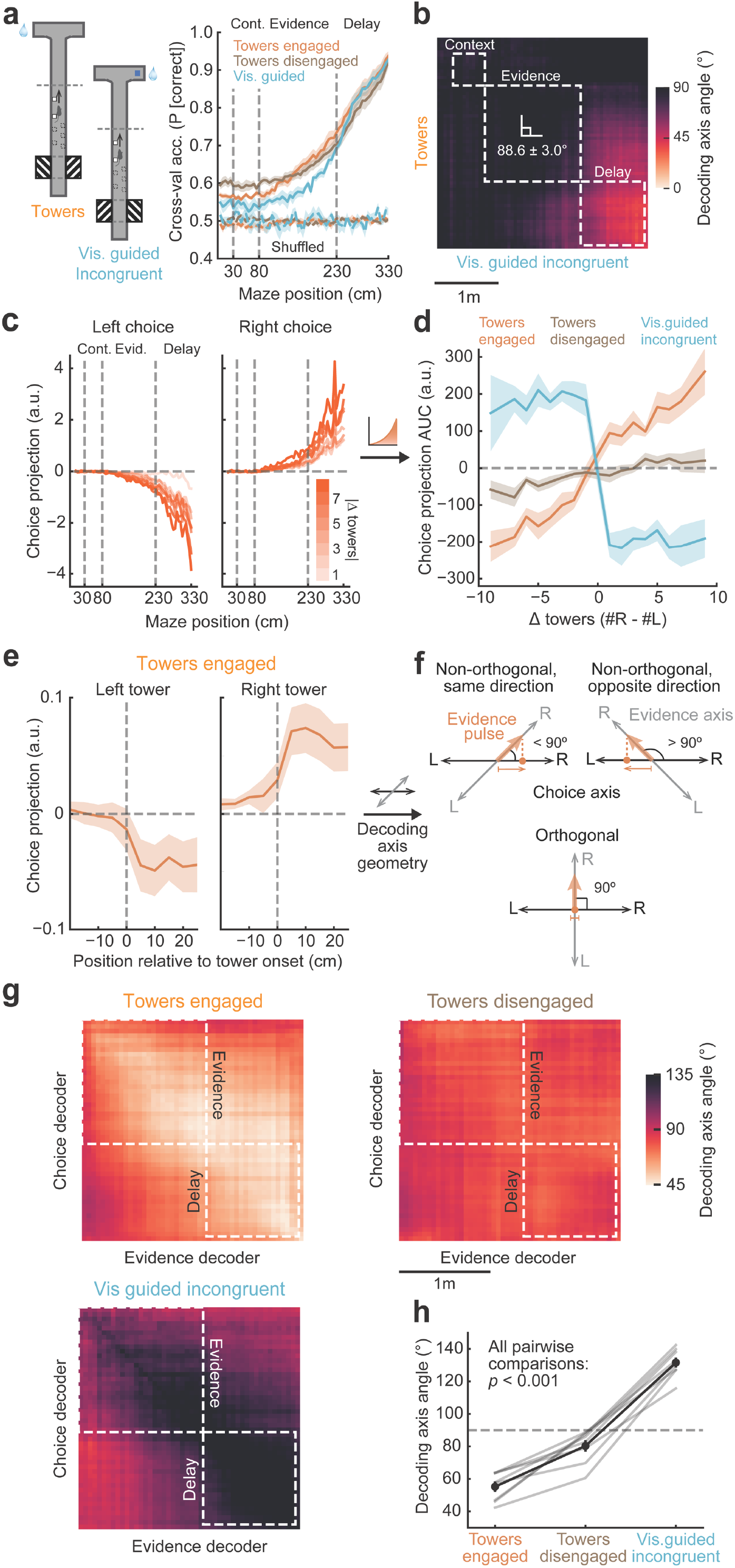
Cholinergic input to the cortex encodes the decision variable in a task- and strategy-dependent fashion. **a**, Accuracy of decoders trained to classify upcoming choice from area-averaged cholinergic-axon activity in different parts of the maze, for engaged- and disengaged-strategy trials in the towers task, and incongruent visually guided trials. Error shades: ± s.e.m. across mice (n = 8). **b**, Angles between the choice-decoding axes for towers and incongruent visually guided trials as a function of maze position, averaged across mice. **c**, Projections of cholinergic activity onto the choice-decoding axes for the towers task (engaged strategy), separately for trials with different amounts of sensory evidence (|Δ towers|, color bar), averaged across mice. **d**, Area under the curves (AUC) in panel c, plotted as a function of sensory evidence for engaged and disengaged strategies in the towers task and incongruent visually guided trials. Error shades: ± s.e.m. across mice (n = 8). Slope of Δ towers × task engagement interaction: *p* < 0.001, slope of Δ towers × task interaction: *p* = 0.025, mixed-effect linear regression. **e**, Projection onto choice-decoding axes aligned to the onset of left and right towers, in the towers task. Error shades: ± s.e.m. across mice (n = 8). **f**, Possible geometric arrangements of choice and evidence encoding. Top left: choice- and evidence-decoding axes forming an acute angle, meaning that evidence pulses drive choice in the same direction. Top right: evidence- and choice-decoding axes forming an obtuse angle, meaning that evidence pulses drive choice in the opposition direction. Bottom: evidence and choice decoding axes are orthogonal, evidence pulses have no influence on choice. **g**, Angles between choice- and evidence-decoding axes across different maze positions, within towers (disengaged and engaged) and incongruent visually guided trials (mouse averages). **h**, Quantification of the average angles in panel g, computed within the evidence region and separately for each mouse (thin gray lines). Black line and error bars: mean ± s.e.m. across mice (n = 8). Print *p*-value is from one-way repeated measure ANOVA and post hoc paired t-tests with FDR correction.

To further understand how sensory evidence contributes to choice-related dynamics in cholinergic axons, we asked how single evidence pulses impacted the decision variable. We aligned single-trial choice-decoder projections to the onset of individual right or left towers. We saw that each tower led to a step-like increase in choice information towards its side (**Fig. 3e**). Although jGCaMP8m has fast dynamics^79^, the exact time course could be impacted by the sensor. Regardless, these findings show that the average evidence ramps in **Fig. 3c** likely correspond to discrete increases locked to the arrival of new evidence. In other words, specifically during the towers task, cholinergic input to the cortex behaves much like a canonical evidence accumulator^80,81^.

As a complementary analysis, we trained linear-regression decoders to predict the amount of sensory evidence from cholinergic activity (**Fig. S6c**). We then examined the relationship between choice and evidence decoding in each task. We reasoned that, for evidence to contribute to choice formation in the towers task, the decoding axes for evidence and choice should have non-zero projections onto each other, i.e., they should be non-orthogonal^82,83^ (**Fig. 3f**). This was indeed the case (**Fig. 3g**). Conversely, compatible with the fact that the evidence from the towers indicates the opposite choice in incongruent visually guided trials, angles in these conditions had opposite directions within the evidence region of the maze (i.e., > 90°). Interestingly, they were approximately orthogonal in disengaged trials in the towers task, when the mice made less evidence-based choices (**Fig. 3h**). Altogether, these analyses show that cholinergic input to the cortex conveys detailed information about cognitive variables in a highly task-specific fashion.

### Cholinergic input to the cortex is selectively required for evidence accumulation

We next asked if task-specific cholinergic input is required for the performance of each task. Specifically, as shown above, this input carries choice information in both tasks, but is evidence dependent only during accumulation. Thus, we reasoned that, if this input is generally required for choice, silencing it should affect performance of both tasks, but if it is specifically required for accumulation, silencing it should only affect towers-task performance. Likewise, in the latter scenario, silencing should specifically impact cortical dynamics supporting evidence accumulation. To test this, we optogenetically silenced cholinergic axons while measuring excitatory cortical activity with mesoscale widefield Ca^2+^ imaging through the cleared skull (**Fig. 4a**). We virally expressed the red-shifted inhibitory opsin Jaws Cre-dependently in basal forebrain cholinergic neurons, in mice that also constitutively expressed GCaMP6s in excitatory neurons (ChAT-IRES-Cre x Thy1-GCaMP6s cross, n = 6, plus one additional ChAT-IRES-Cre mouse without simultaneous imaging, **S7a**). While we imaged cortex-wide activity during task switching, we used a red LED positioned obliquely beneath the imaging objective to illuminate the whole dorsal cortex, thereby suppressing acetylcholine release (note that, although this likely resulted in weaker inhibition than targeting cell bodies in the basal forebrain, it allowed for broader and more specific targeting of axonal input to the cortex). We did this in random subsets of trials during the evidence region of either task. Interestingly, we found that this manipulation selectively and significantly impaired performance of the towers task in experimental animals, but not controls that did not express the opsin (**Figs. 4b, S7b,c**, n = 7 Jaws mice and 23,710 trials, and 6 control mice and 12,826 trials). To quantify which aspects of task performance were affected by cholinergic silencing, we fitted the same behavioral models from **Fig. 1d,e**, separately for ‘LED off’ and ‘LED on’ trials. We found that, while the small impact of trial history on choices was unchanged, the weight of sensory evidence on choice was significantly reduced (**Figs. 4c, S7d**; the effect was roughly even across the evidence region, **Fig. S7e**). In a different set of mice, we systemically administered scopolamine, a pan-muscarinic antagonist, before task-switching sessions (or vehicle as a control, n = 9 mice). Interestingly, this much less specific cholinergic manipulation also resulted in a selective evidence-accumulation deficit (**Fig. S7f**). This suggests that task-specific patterns of cholinergic activity in other regions of the maze (e.g., higher activity in the visually guided task following the evidence region, **Fig. 2d**) are less causal to task performance on a trial-by-trial basis.

**Fig. 4.**
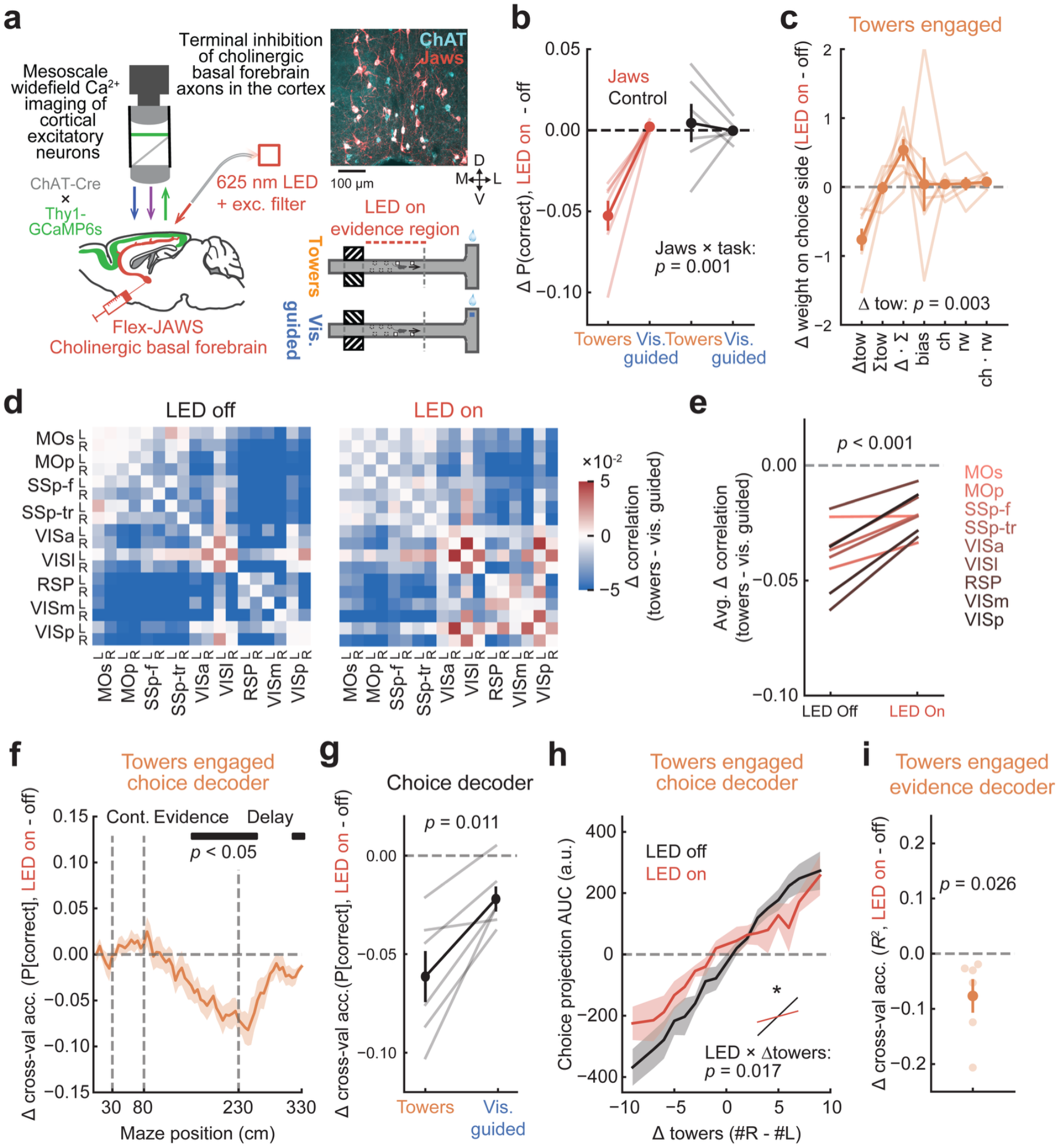
Cholinergic input to the cortex is selectively required for evidence accumulation. **a**, Schematics of simultaneous widefield Ca^2+^ imaging of excitatory neurons and optogenetic silencing of basal forebrain cholinergic axons in the cortex. Image inset exemplifies the specificity of Jaws expression in basal forebrain cholinergic neurons. **b**, Light-induced decrease in performance (‘LED on’ - ‘LED off’) for each task, for experimental mice (n = 7) and control mice not expressing Jaws (n = 6). Error bars: ± s.e.m. across mice. Printed *p*-value is from a two-way mixed ANOVA. **c**, Light-induced decrease in behavioral logistic-regression coefficients (same model as in Fig. 1e) fitted separately to ‘LED off’ and ‘LED on’ towers-task trials in experimental mice (n = 7), showing selective changes in sensitivity to sensory evidence (Δ towers). Error bars: ± s.e.m. across mice. Printed *p*-value is from a paired t-test with FDR correction. **d**, Average task difference in pairwise correlation between excitatory activity of different cortical regions, for control and cholinergic silencing trials. **e**, Average task difference in pairwise correlation per region, i.e., rows of the matrices in panel d. Printed *p*-value is from a paired t-test. **f**, Light-induced decrease in choice-decoding accuracy for the towers task. Error shades: s.e.m. across mice (n = 6). Bar on top indicates maze positions with a significant decrease in accuracy (*p* < 0.05 for at least 3 consecutive positions, t-test). **g**, Average light-induced decrease in choice-decoding accuracy from excitatory activity across maze positions for the towers and visually guided tasks. Error bars: ± s.e.m. across mice (n = 6). Printed *p*-value is from a paired t-test.. **h**, AUC of projection onto choice-decoding axis (as in Fig. 3d) defined by excitatory cortical activity, plotted as a function of sensory evidence for ‘LED off’ and ‘LED on’ trials in the towers task. Printed *p*-value is from a mixed-effect linear regression. **i**, Average light-induced decrease in evidence-decoding accuracy across maze positions for the towers task. Error bars: ± s.e.m. across mice (n = 6). Printed *p*-value is from a one-sided t-test.

Finally, we quantified how task-specific excitatory cortical dynamics were impacted by cholinergic-axon silencing. Like in our previous work^5^, we found that mesoscale activity between cortical regions is less positively correlated during evidence accumulation compared to the visually guided task, compatible with cellular-resolution-imaging studies^14,84^. However, this decorrelation was significantly attenuated during cholinergic silencing (**Figs. 4d,e, S8a**). We next probed how cortical coding of choice and evidence was affected. We separately fitted choice decoders using area-averaged cortical activity during ‘LED off’ and ‘LED on’ trials. Decoding accuracy was significantly reduced with cholinergic silencing, especially during the evidence region (**Fig. 4f, S8b**). This decrease in decoding accuracy was significantly larger than in visually guided-task trials, compatible with the selective behavioral effects in the towers task (**Figs. 4g, S8c**). To understand how the disruption of choice by cholinergic silencing related to sensory evidence, we again projected activity onto the choice-decoding axes, separately for different amounts of evidence. Like in cholinergic axons, choice information at the level of large-scale cortical excitatory dynamics was linearly related to evidence strength. Cholinergic silencing impaired this relationship, significantly decreasing the slope of the choice-information curve (**Fig. 4h**). The accuracy of evidence decoders was also significantly decreased during cholinergic silencing in the towers task (**Figs. 4i, S8d**). This was even though sensory responses to individual towers were unaffected (**S8e**). Thus, compatible with the cholinergic-axon imaging and behavioral deficits from silencing, cholinergic input appears specifically necessary for the emergence of choice and evidence signals across the cortex during accumulation.

## Discussion

Using optical measurement and manipulation of cholinergic axons across the cortex during task switching, we have shown that task-specific patterns of cholinergic input orchestrate large-scale cortical dynamics to support specific cognitive processes. In particular, rather than just changing their overall activity levels, cholinergic axons dynamically adjust their task-coding schemes to track the decision variable during perceptual decisions. These results reveal a novel circuit mechanism enabling distributed cognitive computation by cortical circuits and the accumulation of sensory evidence. They also generalize previous findings of specific cholinergic recruitment by sensory modality, task engagement, and spontaneous movement^21,24–26^ to show that, even when movement and sensory input are held constant, acetylcholine release in the cortex directly tracks and shapes cognitive operations on that input.

Our results demonstrate a specific role of cholinergic modulation in the accumulation of sensory evidence, rather than less task-specific cognitive processes. For example, it could be argued that the towers task involves more arousal or attention, processes previously associated with acetylcholine^30,34,35,70^. However, this is unlikely to explain our main findings. Our linear encoding model of cholinergic activity (**Fig. 2**) explicitly accounts for pupil dynamics, a common index of arousal. Moreover, to the extent that attention would track task difficulty, we might expect that choice-related signals would increase with trial difficulty (i.e., decreasing amounts of sensory evidence), which is the opposite of what we observed (**Fig. 3b,c**). Further, during disengaged strategies (which could be associated with less attention to the stimuli), cholinergic axons are still active and track choice, but in an evidence-independent fashion (**Fig. 3a,d**). Finally, the two tasks have different levels of reward expectation, which are signaled by cholinergic neurons^48,49^. In this case, we might expect uniformly higher cholinergic activity during the visually guided task, which is associated with higher reward expectation based on reward rates (**Fig. 1f**). Instead, we observed higher activity when animals accumulated evidence in the towers task.

Thus, cholinergic input to the cortex directly takes part in the accumulation of individually unreliable pieces of evidence. This is partly compatible with a theoretical framework proposing that the cholinergic system is recruited when an animal expects an uncertain environment^72^. However, in that framework, while contextual cholinergic activity shapes cognitive computation, it does not encode task-specific decision variables themselves. Our work should therefore prompt revised theoretical accounts of neuromodulation.

Future work should establish how cholinergic neurons acquire task-specific cognitive signals. Interesting candidates include the projections from frontal cortical regions to the basal forebrain^55,85^. Another important topic for upcoming studies is the microcircuit-level mechanism whereby acetylcholine supports evidence accumulation in the cortex. This neuromodulator acts on different cell types and compartments to favor inter-areal feedforward input propagation over feedback and local recurrent excitation^32,86–88^. Interestingly, feedforward processing has been proposed as a mechanism of circuit-level integration in theoretical work^89^ and signatures of this are seen in cortical recordings during evidence accumulation, notably the progressive increase in integration timescales over successive processing steps^9,90,91^.

Altogether, our results establish the cholinergic basal forebrain as a flexible orchestrator of cortex-wide dynamics and an unexpected, key node of the distributed network supporting evidence accumulation.

## Methods

### Animals

All experimental procedures were approved by Northwestern University’s Institutional Animal Care and Use Committee and were done in accordance with the Guide for the Care and Use of Laboratory Animals. Mice were kept under a 12/12 h reverse light/dark cycle and, with very few exceptions, were group housed. We used a total of 33 mice of various genotypes and both sexes, aged 2–14 months at the time of data collection, as detailed below (note that some animals were used in more than one experiment).

- Behavioral analyses in **Fig. 1**. Crosses between ChAT-IRES-Cre [B6;129S6-*Chat*^*tm2(cre)Lowl*^/J, JAX stock #006410] and flex-jGCaMP8m [*Igs7*^*tm1(tetO-GCaMP8m,CAG-tTA2)Genie*^/J, JAX stock #037718] (n = 10); ChAT-Δneo-Cre [B6;129S-*Chat*^*tm1(cre)Lowl*^/MwarJ, JAX stock #031661] and flex-jGCaMP8m (n = 1); and ChAT-IRES-Cre and Ai96 [B6;129S6-Gt(ROSA)26Sor^tm96(CAG-GCaMP6s)Hze^/J, JAX stock # 028866] (n = 3).
- Cholinergic-axon imaging experiments in **Figs. 2, 3**. Crosses between ChAT-IRES-Cre and flex-jGCaMP8m (n = 8).
- Histology quantification and controls in **Fig. S2**. Crosses between ChAT-IRES-Cre and flex-jGCaMP8m (n = 3), and between ChAT-Δneo-Cre and flex-jGCaMP8m (n = 1).
- Simultaneous optogenetic and widefield imaging experiments in **Figs. 4**. Crosses between ChAT-IRES-Cre and Thy1-GCaMP6s [line GP4.3, C57BL/6J-Tg(Thy1-GCaMP6s)GP4.3Dkim/J, JAX stock # 024275] (n = 6); and ChAT-IRES-Cre and Ai96 (n = 1) for optogenetics without cortical imaging.
- Behavioral controls for optogenetic experiment. Crosses between ChAT-IRES-Cre and Ai96 (n = 4), and ChAT-IRES-Cre and GP4.3 (n = 2).
- Pharmacology experiments in **Fig. S7f**. Crosses between ChAT-IRES-Cre and Ai96 (n = 4), and ChAT-IRES-Cre and GP4.3 (n = 5).

### Surgical procedures and water restriction

Each mouse underwent a single surgical procedure that varied depending on the experiment. For imaging experiments without optogenetics, the procedure was a titanium headplate implant with optical clearing of the intact skull^5,77^. Mice were anesthetized using isoflurane (3% for induction, 1.5% for maintenance) and secured in a stereotaxic frame (Kopf). Following aseptic preparation, hair removal, and subcutaneous administration of local lidocaine (4 mg/kg), the scalp overlying the dorsal skull was excised and the periosteum scraped. Sequential applications of thin cyanoacrylate adhesive layers (Elmers) and clear metabond (Parkell) were then performed. Once cured, the metabond surface was smoothed using a cement polishing kit (Pearson Dental). The headplate was subsequently affixed to the skull with opaque metabond and a thin layer of black or white nail polish was painted over the metabond to completely block off ambient light. Lastly, a uniform thin coat of transparent nail polish (Electron Microscopy Science) was applied across the implant. For pain management, mice received meloxicam at two timepoints (20 mg/kg S.C., perioperatively and 24 hours post-surgery). Warm saline was also administered subcutaneously after the procedure to ensure adequate hydration (0.1–0.3 mL S.C). A homeothermic blanket (Harvard Apparatus) was used to maintain body temperature at 37 °C during the entire procedure.

For experiments involving simultaneous optogenetics and imaging, the mice additionally received viral injections to deliver the inhibitory opsin Jaws to basal forebrain cholinergic neurons (AAV8-flex-Jaws-mCherry, 500 nL/injection, rate of 125 nl/min, titer ∼2 × 10^12^). Pre-surgical preparation happened as above. We then used an air-driven dental drill with a carbide burr to perform four small craniotomies (∼ 500-µm diameter) to bilaterally target two basal forebrain subnuclei, substantia innominata (SI, AP: -0.23 mm from bregma, ML: 1.5 mm, DV: -5 mm) and horizontal limb of the diagonal band (HDB, AP: 0.6 mm, ML: 0.61 mm, DV: -5.4 mm). We then used a microinjection syringe pump (World Precision Instruments) to deliver the virus, and retracted the syringe 5 min after the injection. The craniotomies were then sealed with bone wax (Surgical Specialties) and the cleared skull preparation and headplate attachment proceeded as above. In this case, analgesia consisted of a subcutaneous injection of meloxicam (20 mg/kg) before the procedure and 24h later, as well a single subcutaneous injection Buprenorphine SR (1 mg/kg) before the procedure.

The mice recovered from surgery for at least 5 days before undergoing habituation to experiments, at which point we water restricted them for behavioral training. Daily water allotments of 1–1.5 mL were provided to each mouse based on their *ad libitum* drinking weight, with supplemental fluid administration if body weight dropped below 80% of baseline. Throughout the initial ∼7 days of water restriction, the mice underwent extensive handling and received their water allotment by hand. Behavioral training began once the mice willingly climbed into the experimenters’ hands and displayed no distress behaviors. Following each training session, the mice were given access to an enclosure containing toys and conspecifics for environmental enrichment.

### Histology and immunohistochemistry

We confirmed that jGCaMP8m and Jaws expression was specific to cholinergic neurons with choline acetyltransferase (ChAT) immunostaining in a subset of animals, along with green- and red-fluorophore signal amplification with GFP or RFP staining (n = 3 and 2, respectively). The mice were deeply anesthetized with ketamine (50–100 mg/kg) and xylazine (10 mg/kg) and transcardially perfused with phosphate-buffered saline (PBS, 20 mL), followed by 4% paraformaldehyde (10 mL). The brains were post-fixed in 4% w/v paraformaldehyde for 12–24 h, and then cryoprotected with 30% w/v sucrose. The tissue was sectioned coronally in 50-µm slices with a cryostat (Leica). The slices were washed with PBST (0.3% Triton-X in PBS, 3 × 10 min), incubated with blocking buffer for 2h at room temperature (6% v/v normal donkey serum), and incubated for two nights with primary antibody at 4 °C. This was followed by another PBST wash (3 × 10 min), 2 h of incubation with secondary antibody at room temperature, a final PBST wash (3 × 10 min), and mounting on glass slides with Fluoromount-G™ with DAPI. We used the following antibodies:

- ChAT staining. Primary: goat anti-ChAT polyclonal, 1:75 (Millipore, AB144P). Secondary: donkey anti-goat Alexa Fluor 647, 1:500 (Invitrogen, A32849).
- GFP (GCaMP). Primary: chicken anti-GFP polyclonal, 1:2000 (Abcam, AB13970). Secondary: donkey anti-chicken Alexa Fluor 488 IgY, 1:500 (Jackson ImmunoResearch, 703-545-155).
- RFP (mCherry). Primary: rabbit anti-RFP polyclonal, 1:200 (Rockland #600-401-379). Secondary: donkey anti-rabbit Alexa Fluor 594, 1:500 (Invitrogen, A21207).

To quantify expression overlap, we acquired z-stacks from across the basal forebrain with 20× magnification using a confocal microscope (Nikon W1 Spinning Disk). Cell counts were performed manually using ImageJ and averaged across fields of view. Finally, we also quantified jGCaMP8m expression in cortical interneurons in ChAT-IRES-Cre- and ChAT-Δneo-Cre-based crosses, using GFP-based amplification and manual counting as above. In this case, images from across the cortex were acquired with 4× magnification using a widefield epifluorescence microscope (Nikon Ti2 Widefield). In the latter case, cortical regions of interest were manually drawn for the quantification of cell densities.

### Behavioral paradigm

#### Virtual-reality (VR) apparatus

We used a custom-built apparatus described in detail elsewhere^77^. Briefly, head-fixed mice ran on a Styrofoam© ball suspended by compressed air. Ball displacements were measured using an optical velocity sensor lying underneath it and transformed into x-y and angular movements in the virtual world, in closed loop, using custom code written using ViRMEn^92^, running on a PC. The virtual world was projected onto a spherical screen (17” internal diameter) using a DLP projector with 120-Hz refresh rate, image resolution of 768 × 1024 pixels and RGB color balance of [0 0.4 0.5]. Reward was delivered through a metal spout by opening a solenoid valve controlled by a TTL from a DAQ card. The apparatus was enclosed in a custom-designed sound-attenuated cabinet.

#### Task design

Mice switched between two tasks modified from our previously published designs, towers and visually guided^5,61^. Both tasks happened within the same virtual T-maze with the stem measuring 10 × 330 × 5 cm and side arms measuring 25.5 × 11 × 5 (width × length × height, referred to as x × y × z). The maze was divided into the following regions along the y dimension: start box (30 cm), context region (50 cm), evidence region (150 cm), delay region (100 cm), and arm (20 cm). Other than the context region, the maze was decorated with wall paper design to provide optical flow. At the beginning of each trial, the mouse was teleported to the start box and, as it entered the context region, bilaterally symmetrical, full-contrast gratings of oblique orientation covering maze walls were revealed. A 155° grating indicated that the upcoming trial was visually guided, while 25° indicated towers task. For a subset of animals in the optogenetics experiments, context was instead indicated by wallpapers of different color (n = 8, both task variants: n = 1; blue for visually guided and yellow for towers). In the towers task, the mouse then entered the evidence region, where it experienced briefly flashing white towers on either side of the maze (2 × 2 × 6 cm). Tower counts and positions followed spatial Poisson processes with different rates on the rewarded and unrewarded sides (3.7 and 1.1 towers/m, respectively, with a refractory period of 12 cm and maximum density of 4.8 towers/m). These were drawn on each using an algorithm we described elsewhere^61^. Towers appeared when the mouse was 10 cm away from them along the y dimension, and disappeared 200 ms later. This resulted in a median tower count of 5 and 1 on the rewarded and non-rewarded sides, with a respective range of 1–10 and 0–8. The delay region did not contain towers. After it, the mouse was rewarded with a drop of 10% v/v sweet-condensed milk (4–8 µL) for turning towards the side with the highest tower count. Errors were indicated with a sound. The maze froze for 1 s after the end of the trial and the screen went black for an inter-trial interval of 2 s or 10 s, respectively after a correct choice and an error. The visually guided task also contained the towers following the same generative statistics and with the same overall timing structure. Additionally, a tall blue tower (the visual guide, 2 × 2 × 50 cm) placed on the rewarded side arm was permanently visible from right after the mouse entered the evidence region to the end of the trial. The visual guide was on the side opposite the one with the majority of towers on 20% of the trials. In both tasks, whether the rewarded side was left or right was drawn from a dynamically varying probability to counteract biases measured over the session, following an algorithm we described previously^61^. Task switches occurred stochastically with an approximately flat hazard rate. Trial-block lengths were drawn from a geometric distribution with *p* = 0.35 and a range of [8 30] trials for the towers task and [5 15] trials for the visually guided one. The overall longer blocks for the towers task were chosen to discourage the mice from only engaging with the visually guided task to retrieve most of their reward and to yield roughly equal engaged-trial counts for each task, based on pilot studies (we define engaged based on GLM-HMMs, see *Behavioral Analysis* below). Behavioral sessions had a median trial count of 241, median trial duration of 9.9 s, median of 30 task switches, and median block length of 9 trials for the towers and 5 for the visually guided task.

#### Training procedure

After extensive habituation (see above), the animals underwent initial motor training to control the treadmill by running on a 50-cm-long linear track. In this pre-training phase, mice were rewarded when they reached the end of the track in less than 60 s. Mice remained in pre-training until they could reliably obtain ∼1 mL reward in 20–30 min, which took a median of 4 sessions. They then underwent a software-automated shaping pipeline of 11 steps, similar to the one we described previously^61^, and with the introduction of the task-switching contingency in the last step. Details about the steps and criteria are provided in **Tables S2–S4**.

### Mesoscale widefield Ca^2+^ imaging

We used our custom widefield macroscope for imaging, described in detail elsewhere^77^. Briefly, it consisted of 1×-0.63× tandem lenses (Leica) with a dichroic beamsplitter and green emission filter (Semrock, FF495-DI03-50X70 and FF01-525/45), a Prime BSI Express sCMOS (Teledyne), and alternating illumination with blue and violet LEDs (Cairn, 470 nm and 405 nm respectively, 40-nm bandpass filtered, ∼2 mW/cm^2^ power at the sample). Images were captured at 20 Hz with 1024 × 1024 pixel resolution (6.8 µm pixel size, approximately 7 × 7 mm field of view), controlled by µManager on a PC. We synchronized imaging with behavioral data by routing sCMOS exposure signals through a National Instruments DAQ card to the VR PC, integrating the signals directly into the behavioral data file.

### Eye tracking

We performed video-based eye tracking. To avoid obstructing the animal’s view of the VR environment, we positioned a ∼4-cm^2^ mirror (First Surface Mirrors) at an angle under the mouse’s left eye and below ball level, and filmed the mirror using a camera system positioned above the mouse, ∼25 cm from the mirror. We used a CMOS blackfly camera (Teledyne, 1920 x 1200 pixels) with a 35-mm focal-length lens (Tamron M112FM). For illumination, we used an infra-red LED with a broadband diffuser (740 nm, Digikey), driven by a custom-printed circuit board and placed ∼10 cm behind the mirror so as to form a ∼90° angle with the camera. Images were acquired at 15 Hz using a second instance of µManager running on the same PC as the widefield camera, and synchronized by saving exposure signals into the behavioral data file as above.

### Optogenetics and systemic pharmacology

In experiments using simultaneous imaging and optogenetic silencing of cholinergic axons, we used a 620-nm LED (Thorlabs Solis, DC22 driver, 10 mW/cm^2^), directed to the sample in dark-field illumination mode using a ½” liquid light guide (Thorlabs) attached underneath the objective at a ∼45° angle using a custom 3D-printed shield that also protected the objective from ambient light. LED light was pulsed as a 10-Hz square wave with a 50% duty cycle within the evidence region of the maze in each task. LED-on trials were randomly drawn from a binomial distribution with *P* = 0.33. The LED was controlled by the VR software using TTL pulses.

In experiments involving cholinergic pharmacology, the mice were injected with scopolamine (1 mg/kg in PBS) or vehicle (PBS) intraperitoneally, 15 min before the start of the behavioral session. Each animal underwent 2–8 scopolamine and vehicle sessions, with injection order determined randomly.

### Behavioral analysis

#### GLM-HMM and behavioral-data selection

To select trials where the animals were employing the intended behavioral strategies, we fitted Bernoulli generalized linear models combined with a hidden Markov model (GLM-HMM)^67^ to session-concatenated data, separately for each animal and each task. Briefly, the GLM-HMM is composed of a hidden Markov model with *K* states and *K* state-specific Bernoulli GLMs. The transitions between the states are governed by a *K* × *K* transition matrix and follow the Markov property, such that the state on a given trial only depends on the previous trial. For both tasks, the Bernoulli GLM is formulated as:

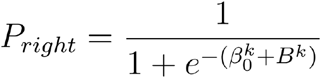

where 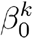 is the state-specific intercept (i.e., side bias) for state *k* and *B*^*k*^ is the linear combination of state-specific predictors for state *k* and for each task. For the towers task:

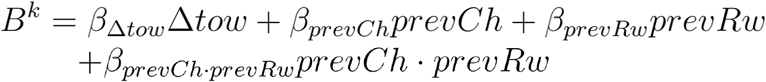

where Δ*tow* is the difference between right and left tower count, *prevCh* and *prevRw* are single indicator functions for chosen sides and whether the animal was rewarded in the previous trial, and *prevCh* · *prevRw* is the interaction between *prevCh* and *prevRw* (i.e., a win-stay/lose-switch term). For the visually guided task:

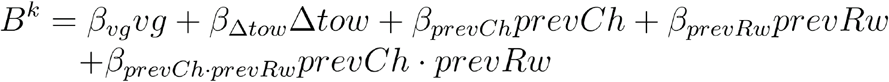

where an additional predictor *vg* indicating the side of the visual guide is included. The GLM-HMM models were fitted using the Expectation-Maximization algorithm for a maximum of 1000 iterations. The fitting process was repeated 10 times with different initial values and the fitted model with the highest log likelihood was chosen. After fitting, we designated the states as ‘engaged’ and ‘disengaged’ by two thresholding criteria. For a state to be labelled as ‘engaged’, 1) the minimum average behavioral accuracy of the state must be greater than or equal to 0.6 and 0.9 for the towers and visually guided task respectively; and 2) the average accuracy of the state must be greater than its side bias (computed as the difference between right and left trial accuracies). States failing to meet either of these criteria were labelled as ‘disengaged’. We set *K* = 5 and 3 states for the towers and visually guided tasks, respectively. Note that, because the primary objective of the GLM-HMM model fitting is to select trials where the animals were employing the intended behavioral strategy, we allowed for a slightly high number of states for the towers task to minimize artificial merging of high- and low-performance states (based on pilot modeling). Trial selection with GLM-HMM model fitting resulted in a 44.3% (18,969-trial) and 84.6% (21,771-trial) yield for engaged states in the tower and visually guided tasks, respectively, in the behavioral and widefield cholinergic axon imaging dataset (**Figs. 1–3**). For the optogenetic perturbation and widefield imaging dataset (**Fig. 4**), 66.0% (13,324 trials) and 85.6% (12,132 trials) of the towers and visually guided trials were assigned to engaged states.

#### Psychometrics and overall performance

Overall performance was defined as the fraction of trials in which the mouse turned to the same side as the majority towers in the towers task and as the visual guide in the visually guided task. For psychometrics, trials were binned by Δ towers between -10 and 10, with bins of 1 at [-7 7] and a single bin at [-10 -7) and (7 10] to ensure comparable occupancy in the edge bins (for which trials are rarer). The fraction of trials in which the mice turned right were computed separately for each bin and each mouse. The population means and standard errors were then computed over mice. A sigmoidal function was then fit to both the within-subject and population averages:

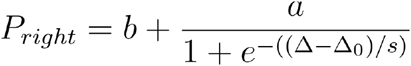

where *b* is a symmetrical offset (lapse rate), *a* is the amplitude of the curve (distance between asymptotes), Δ is the difference between right and left tower counts (the independent variable), Δ_0_ is the horizontal offset of the curve, and *s* controls the slope of the curve.

#### Logistic regressions (Bernoulli GLMs)

We fitted mixed-effects Bernoulli GLMs to predict the probability of a rightward choice from one of two sets of predictors. In both cases the models were formulated as:

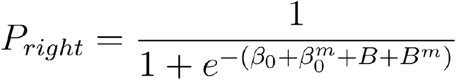

Where *β*_0_ is a general offset (i.e., side bias) term, *B* is a model-specific set of predictors as described below, and 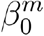 and *B*^*m*^ are the random-effects terms that capture variability across mice *m*, respectively for offset (side bias) and each term *i* out of *I* inside of *B*. 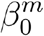 and *B*^*m*^ are modeled together as a zero-mean multivariate Gaussian:

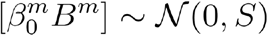

where *S* is an *n* × *n* covariance matrix, with *n* = *I* + 1 and diagonal elements being predictor-level variances, 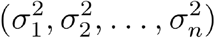.

One of the models (**Figs. 1e, 4c, S7d**) contained terms related to both evidence and trial history:

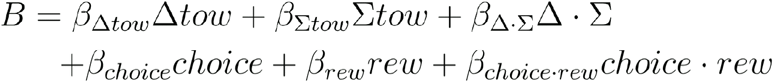

Where Δ*tow* is the difference in right and left tower counts, ∑*tow* is the total number of towers (sum of left and right tower counts), Δ · ∑ is the interaction between Δ*tow* and ∑*tow, choice* is an indicator variable for the chosen side in the previous trial, *rew* is an indicator variable for whether or not the previous trial was rewarded, and *choice* · *rew* is the interaction between *choice* and *rew* (i.e., a win-stay/lose-switch term).

The other model (**Figs. 1d, S7e**) was used to estimate how the mice weighed evidence from different parts of the maze (akin to a psychophysical kernel):

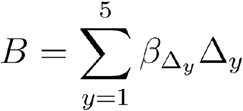

Where *y* are equally spaced y position bins within the evidence region of the maze (excluding the first 10 cm, since 10 cm is the first possible tower position) and Δ_*y*_ is the difference in towers counts on each side for that maze segment.

#### Pupil tracking

To track the mouse’s pupil dynamics, we first converted raw tif image sequences to mp4 videos using ffmpeg. Using the DeepLabCut (DLC) GUI, we extracted a total of 80 frames from 4 behavioral sessions with its k-means-based frame selection feature and manually labeled 8 pupil coordinates uniformly spaced around the edge of the pupil^93^. We then trained a supervised, markerless pose-estimation model using Lightning Pose to track pupil-marker coordinates in all session videos^94^. The Lightning Pose model used a video augmentation pipeline implemented in the DLC 2.0 package and a resnet50_animal_ap10k model backbone. We used a mean-squared-error loss, computed from heatmap regression of the 2D Gaussian kernel of the manually labels and the model-predicted probability heatmap of the location of each marker on the frames. The model was trained for 750 epochs with early stopping using Adam optimizer, an initial learning rate of 1e-3 and a multi-step learning rate scheduler from PyTorch decaying by a factor of 0.5 in epochs 150, 200, 250. From the model output, an ellipse was subsequently fitted to these coordinates (EllipseModel, scikit-image), and pupil area was computed in squared pixels from the fitted major and minor axes. Example frames of each session were manually inspected for general quality control of model-predicted locations of the coordinates. Pupil area was further filtered by 1) excluding frames with a marker-averaged likelihood below 0.98 (estimated by the Lightning Pose algorithm) to exclude frames where the pupil was occluded; and 2) allowing for a maximal single-frame decrease of 20% from the session median computed over frames that satisfied criterion 1, to distinguish squinting or blinking of the eye from pupil constriction. Frames excluded with these criteria were assigned NaN values.

#### Quantification of pupil diameter and running patterns

Trial-by-trial pupil area and running speed were first binned to 5 cm bins. Trial data were combined across sessions for each animal to compute a within-subject average, and the population mean and standard errors were computed over the within-subject averages. 11 of the 14 mice used in **Fig. 1** had simultaneous pupil tracking (26,478 engaged trials). For the analysis in **Fig S2d–e**, we similarly computed the within-subject averages and then population mean and standard error but separately for the first five trials of a task block.

#### Optogenetics

For the analysis of optogenetic-silencing effects on behavior, we computed fraction correct, psychometrics and logistic regression as above, separately for ‘LED on’ and ‘LED off’ trials. We then performed paired statistical tests as described below (*General statistics*). For the mixed-effects models, paired statistics were performed on the random-effect terms.

### Widefield preprocessing

We have described preprocessing procedures in detail elsewhere^77^. Briefly, we performed motion correction using phase correlation and spatially binned frames to 128 × 128 pixels (pixel size of 54.4 µm). We then computed separate pixel-wise Δ*F/F* for each excitation wavelength, Δ*F/F*_*B*_ and Δ*F/F*_*V*_, with baseline defined as the mode of each Δ*F/F* over a 60-s running window. We then applied linear regression to scale Δ*F/F*_*V*_ to the troughs of Δ*F/F*_*B*_, followed by smoothing with a 400-ms Gaussian kernel, thereby obtaining traces denoted as 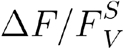 . Δ*F/F* was then computed as a divisive correction, 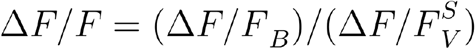 . Finally, we masked large blood vessels using an automatic detection algorithm computed on the unbinned average frame as follows. We first used adaptive median filtering to mask any pixels deviating from the average intensity of a smoothed mean image by 1.6 *σ* in the negative direction or 10 *σ* in the positive direction. We combined this with a second mask computed with the same method, but on the first principal component of the average raw-fluorescence image time series instead of the average frame, and with a manual mask to block off-brain pixels (e.g., headplate) and the sagittal sinus. Finally, pixels were grouped data into anatomically defined cortical areas by registering the average image to the Allen Brain Atlas ccf3.0 using an affine transformation.

For experiments with simultaneous optogenetics, we corrected the signals for residual contamination by the red light. We took the difference between the average raw fluorescence intensity of the three frames before and after the optogenetic LED onset/offset separately for blue and violet excitations. The differences across all the LED-on trials of a session were averaged to estimate the amount of residual contamination for that session. The estimated contamination was then subtracted from both the blue- and violet-excited raw fluorescence.

### Linear encoding model of neural activity

We fitted a mixed-effects linear encoding model to area-averaged cholinergic-axon or excitatory-neuron imaging data, with L2 regularization (**Figs. 2g,h, S5, S8e**). A single model was fitted in the time domain (rather than maze space) for z-scored Δ*F/F* concatenated across behavioral sessions, only containing data points between the start and the end of the delay regions of the maze in both tasks, and only for trials from task-engaged strategies as determined by GLM-HMMs fit to behavioral data (see above). The models were fitted with 5-fold cross validation, with splits at the level of whole trials to prevent inflation of goodness-of-fit estimates because of temporal autocorrelations introduced by the Ca^2+^ sensor (i.e., it could be trivial to explain variance in temporally adjacent data points in training and test sets). The data splits were also done hierarchically over mice and sessions to prevent the overrepresentation of individual mice and experiments. The model was parameterized as follows:

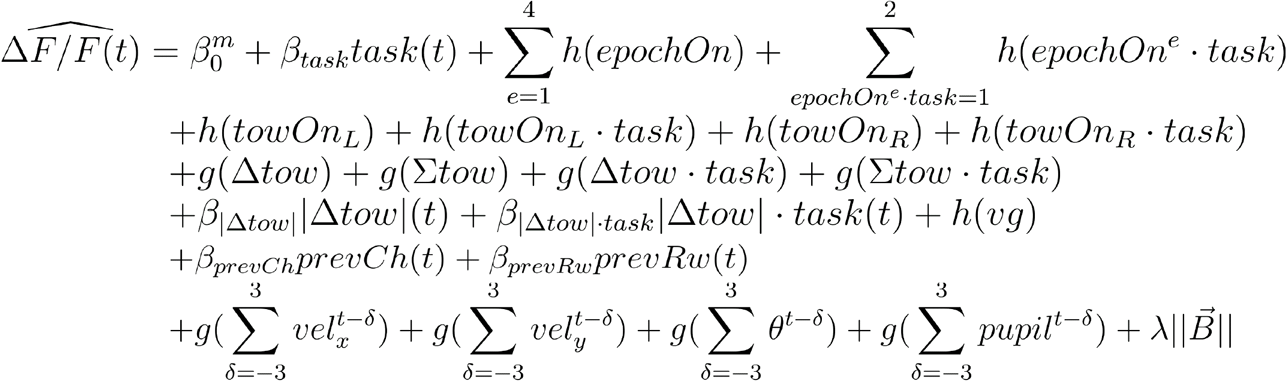

Where 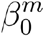, the mixed-effects term, is a random offset for each mouse, modeled as a multivariate Gaussian as described above, *g*(*x*) is a polynomial expansion of the predictor *x*(*t*) up to the 3^rd^ degree, with degree 0 corresponding to a step function:

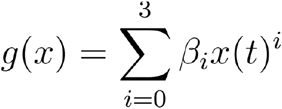

*h*(*x*) is a convolution of *x*(*t*) with a 3^rd^-degree basis set of 7 splines 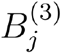 over 2 seconds:

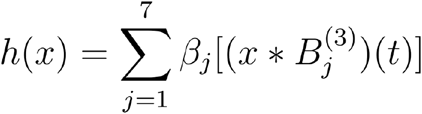

And *task* is an indicator variable for current task being performed, *epochOn*^*e*^ are delta functions at the onset of each task epoch (by convention, *e* = 1 is context, *e* = 2 is delay, *e* = 3 is evidence, *e* = 4 is start box); *towOn*_*L(R)*_ is a delta function at the onset of each left (right) tower; Δ*tow* is the cumulative amount of sensory evidence (#R - #L tower counts); |Δ*tow*| is the absolute of that; ∑*tow* is the cumulative total tower count (#R + #L); *vg* is the side of the visual guide (-1 or 1, or 0 in the towers task); *prevCh* and *prevRw* are single indicator functions for chosen sides and whether the animal was rewarded in the previous trial, present throughout the current trial; *vel*_*x*_, *vel*_*y*_ and *θ* are respectively instantaneous x and y velocities and virtual view angle; *pupil* is pupil area. *vel*_*x*_, *vel*_*y*_, *θ* and *pupil* were all additionally shifted by ± 3 time points, with missing data points introduced by time-shifting filled by linearly extrapolating from 5 (*vel*_*x*_, *vel*_*x*_, *θ*) and 3 (*pupil*) timepoints before for the end of a trial and after for the beginning of a trial. Finally,·*task* denotes a task interaction term; 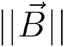 is the L2-norm of the coefficient vector and *λ* is the regularization parameter, determined by cross-validation.

For the reduced model analysis, we dropped entire groups of predictors and refitted the model. For instance, the model without tower onsets dropped the entire spline basis set. We then quantified the contributions by taking the difference between the cross-validated *R*^2^ (computed in the test sets) of the full and reduced model from each cross-validation fold, averaged across hemispheres for each area. To test for statistical significance, the *R*^*2*^ difference of a given area is tested against an *H*_0_ = 0 with a one-sample t-test, and FDR correction was used to correct for multiple comparisons across areas.

### Decoding models

#### Model training

We used logistic regression to decode binary variables (choice, task, behavioral state, trial outcome) and linear regression for evidence (**Figs. 3, 4f–i, S4c,d, S6, S8b–d**). For each animal, activity was averaged within anatomically defined regions and concatenated across sessions, and a single model was fit per animal. Model quantification and statistics were then performed across animals. The models were fitted with L2 regularization and 5-fold cross-validation (with the optimal regularization determined with cross validation). For all decoders, separate models were trained per binned maze position (bin size of 5 cm). For the decoding of binary variables, the decoders were fitted with class weights inversely proportional to class proportions in the training data, and the performance of a decoder was assessed by its balanced accuracy ([sensitivity + specificity] / 2) computed on held out data. For decoding evidence, the coefficient of determination (*R*^2^) was computed on held out data. To obtain shuffled decoder performance, we randomly shuffled the trial labels of the entire dataset and the regularization strength of a given shuffled decoder was chosen to match the best-performing decoder for the same position on non-shuffled data. Statistical significance of decoder performance compared to shuffled data was assessed using a two-sided independent t-test. A given position must have a *p* value < 0.05 and in the neighborhood of at least three consecutively significant positions to be considered statistically significant.

#### Decoding axes and choice projections

Decoding axes were defined as the vectors of decoding weights for the optimal model. Cross-decoder angles were computed as the arccosine of the dot product between the unit vectors of decoding axes. For analyses involving projections onto the decoding axes, the projections were computed on held-out data in each cross validation fold. To compute the area under the curve (AUC) of the choice projections, the pre-evidence projection baseline of each trial, computed as the mean over the start and context region, was first subtracted. The area under the baseline-subtracted projection in the evidence and delay regions was then numerically integrated using the composite trapezoidal rule. To compute choice projections with view angles regressed out, the neural activity of each brain area, on each trial, and in each maze position bin, was first regressed against the view angle at that position on the same trial. After this, the model residuals were used to train the choice decoder as above.

#### Evidence-triggered average of choice projections

For each trial, we first computed vectors indicating in which maze position bins tower onsets took place for left and right towers. After fitting the choice decoders for each maze position bin, we computed the projections onto the decoding axes across the maze position bins for each trial. Projections around the position where a tower onset took place (20cm before, 25 cm after the onset position) were then averaged within animals, from which the mean and s.e.m. of the evidence-triggered choice projection profile was computed.

### Inter-area correlation analysis

For the analysis in **Fig. 4d,e**, for each trial we averaged *ΔF/F* within anatomically defined regions and computed the pairwise linear correlation coefficient between areas for the time points during which the animal was within the evidence region of the maze. Correlation coefficients were then averaged across trials from each task and subtracted.

### General statistics

We detail shuffle-based tests in their respective sections. For parametric analyses, we used t-tests for single comparisons (paired where appropriate). For comparisons between multiple groups, we used analysis of variance (ANOVA, one- or two-way, with or without repeated measures, or mixed). Whenever multiple comparisons were involved, we either performed FDR correction^95^ (e.g., multiple brain areas, post hoc pairwise comparisons), or we set a consecutive threshold of 3 when temporal correlation between data points were expected^96^ (e.g., decoder performance across positions).

## Data availability

The full dataset will be publicly released upon publication of this work.

## Code availability

The source code will be publicly released upon publication of this work.

## Acknowledgements

This work was supported by the US National Institutes of Health (NIH) grant R00MH120047, US National Science Foundation grant IOS-2337351, Simons Foundation grants 872599SPI, NC-GB-PilotExt-00002091, Alfred P. Sloan Foundation grant SP-2022-19027, Scialog grant SA-MBC-2023-079a from the Research Corporation for Science Advancement (all to L.P.), as well as an NIH T32 to J.T. (parent award 5T32MH067564). We thank the GENIE Project/Janelia for sharing the jGCaMP8m mice. We thank Julia Cox, Tobias Donner, and Alessandro Toso for thoughtful comments on this manuscript, and members of the Pinto lab for helpful discussions.

## Author contributions

L.P. and J.K.L. designed the research. LP., J.K.L., P.S.S. and A.P.R designed the behavioral paradigm. J.K.L. collected the widefield and optogenetics data, with help from E.M.. J.T. implemented eye tracking and oversaw the histology experiments. J.T. and E.M. performed the histology. E.M. trained the mice. J.K.L. analyzed the data. L.P. and J.T. secured the funding. L.P. and J.K.L. wrote the paper. L.P. supervised the work.

## Supplementary Figures and Tables

**Fig. S1.**
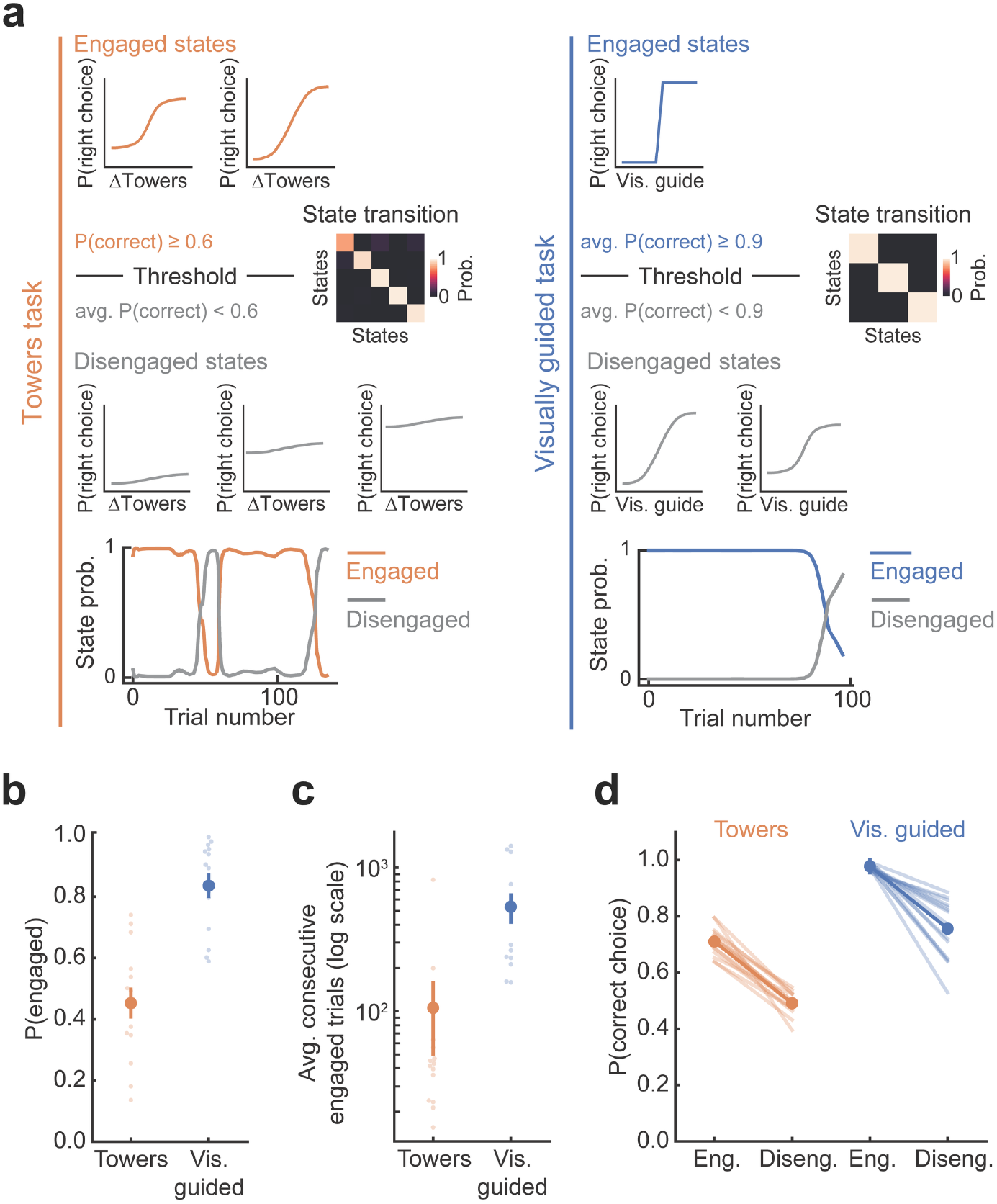
Quantification of behavioral strategy and trial selection with GLM-HMMs. **a**, Single-mouse examples of GLM-HMMs, fitted separately for each task. Top: psychometric curves for engaged states, determined based on the overall P(correct) threshold printed underneath. Right: HMM transition probability matrix. Middle: same as top, for disengaged states. Data from all engaged (disengaged) states were combined for analysis and are collectively referred to as engaged (disengaged) strategies. Bottom: Example session showing the posterior probability of each strategy as a function of trial count. **b**, Probability of engaged strategies across trials for each task. Faint data points: individual mice (n = 14), saturated-color circles and error bars: mean ± s.e.m. across mice. **c**, Number of consecutive trials in an engaged-strategy state. Conventions as in b. d, Overall performance in engaged and disengaged strategies. Faint lines: individual mice (n = 14). Saturated-color data points, lines, and error bars: mean ± s.e.m. across mice.

**Fig. S2.**
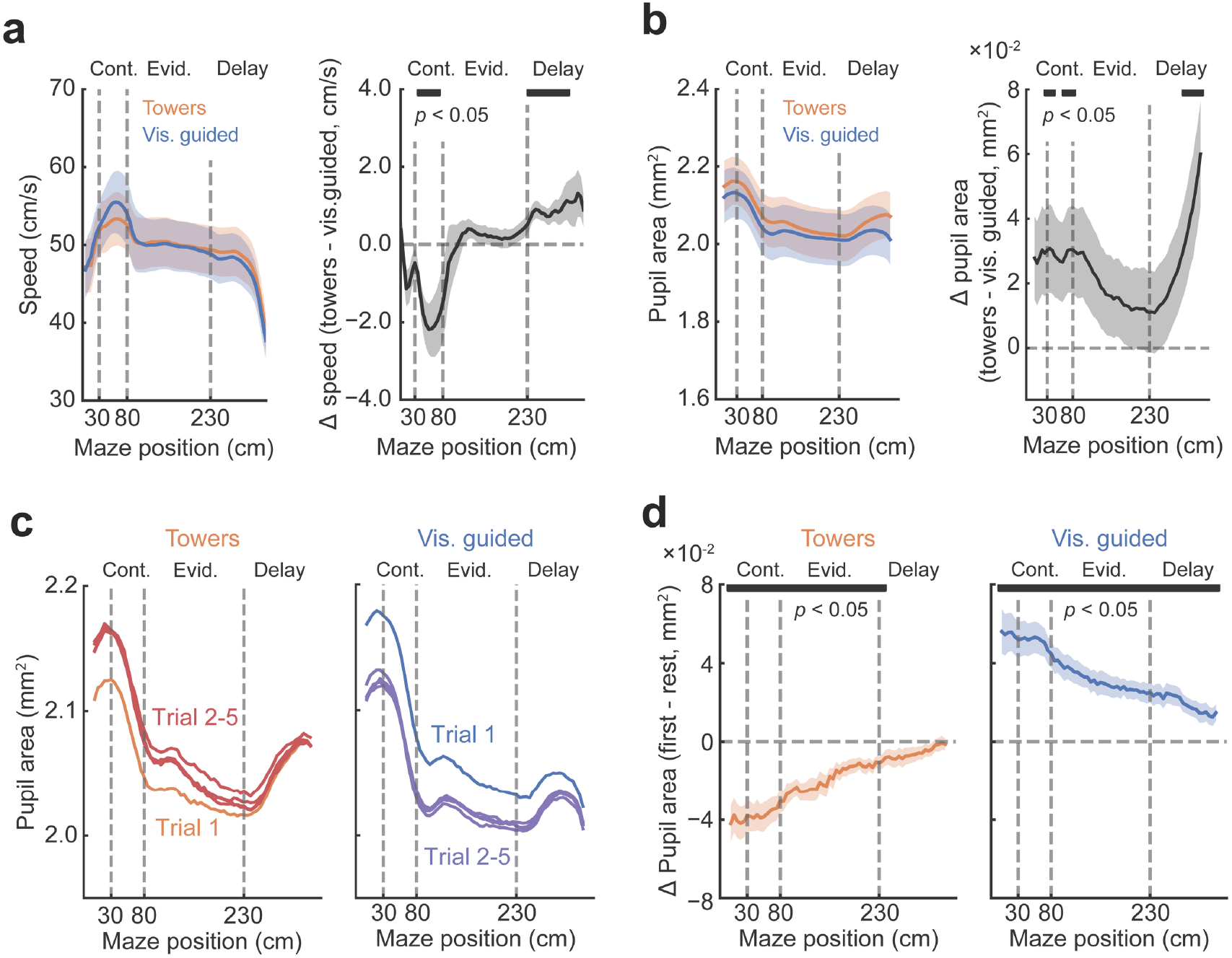
Pupil dynamics and running patterns during task switching. **a**, Left: average running speed as a function of maze position for each task. Right: average task difference in running speed as a function of maze position. Bar on top indicates maze positions with a significant task difference (*p* < 0.05 vs. zero for at least 3 consecutive positions, t-test). Error shades for both: s.e.m. across mice (n = 14). **b**, Same as a, for pupil area (n = 11). **c**, Average pupil area as a function of maze position, plotted separately for the first five trials after a switch into the task indicated on top. Error estimates are omitted for clarity. Note that trial 1 in the towers task resembles more trials 2–5 of the visually guided task (i.e., the previous block), and vice versa. **d**, Average difference in pupil area between the first and subsequent trials as a function of maze position, quantifying the effects in c. Note that pupil dynamics progressively approach its average task patterns over the course of the first trial. Error shades for both: s.e.m. across mice (n = 11). Bar on top indicates maze positions with a significant task difference (*p* < 0.05 vs. zero for at least 3 consecutive positions, t-test).

**Fig. S3.**
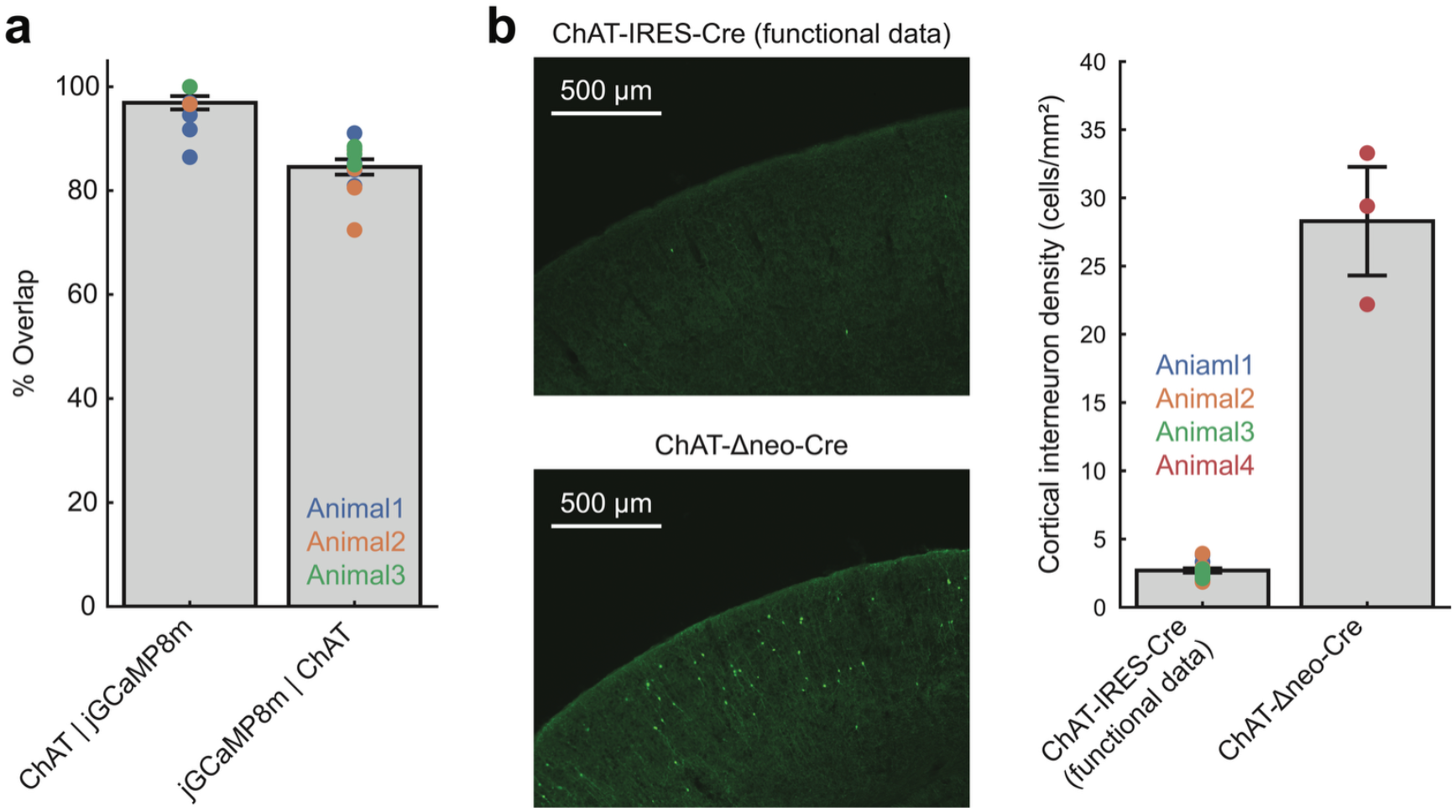
Validation of transgenic strategy for cholinergic-axon imaging. **a**, Quantification of the specificity (ChAT | jGCaMP8) and penetrance (jGCaMP8 | ChAT) of transgenic jGCaMP8m expression in basal forebrain cholinergic neurons. Data points correspond to individual fields of view, colored according to the animal they came from. Error bars: ± s.e.m. across fields of view. **b**, Left: Example coronal fields of view of the cortex from two ChAT-Cre lines crossed with flex-jGCaMP8m, showing minimal expression in cholinergic interneurons in ChAT-IRES-Cre (the one used for imaging experiments) and denser expression in the Δneo line (not used for experiments because of this reason). Right: density of jGCaMP8m-expressing cortical interneurons in the two lines. Conventions as in a. Note that, assuming isotropism, the interneuron density in the ChAT-IRES-Cre line corresponds to an average of ∼0.01 cortical interneurons per widefield pixel, meaning that we are almost exclusively imaging from basal forebrain cholinergic axons.

**Fig. S4.**
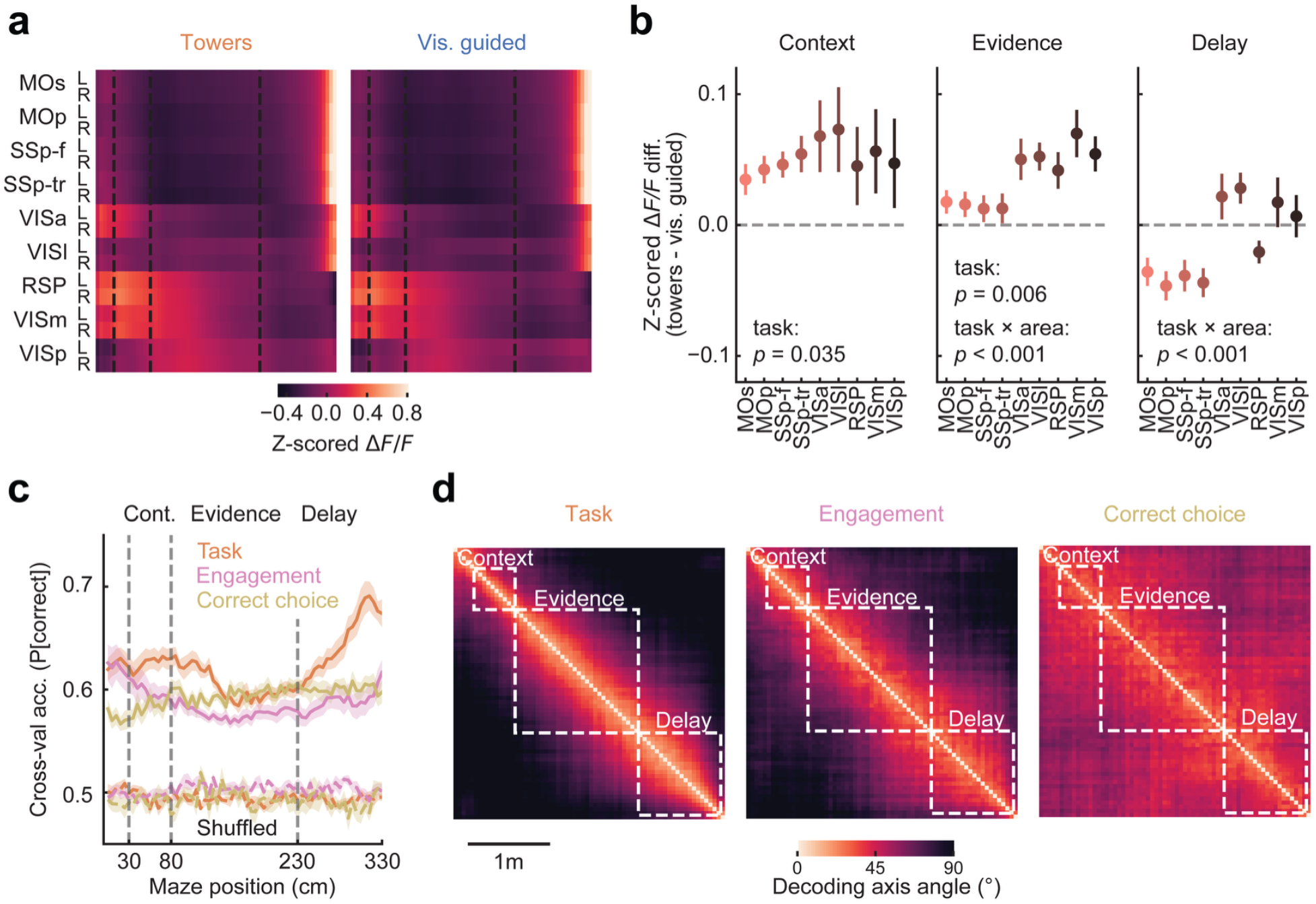
Cholinergic input to the cortex multiplexes cognitive variables. **a**, Grand average of activity in each task, across maze positions (columns), for cholinergic axons innervating different anatomically defined cortical regions (rows) (n = 8 mice, 21,044 trials). **b**, Difference in activity between tasks, averaged by maze region. Error bars: mean + s.e.m. across mice (n = 8). Printed *p*-values are from a two-way repeated measures ANOVA. **c**, Average cross-validated accuracy of decoders as in Fig. 2e, restricted to trials following a reward to control for overall differences in task performance. The decoders were trained to classify the current task, engaged vs. disengaged strategy in the towers task, or whether the upcoming choice was correct in the towers task, from area-averaged cholinergic-axon activity at different maze positions. Shaded regions: ± s.e.m. (n = 8 mice). All positions have significantly higher decoding accuracy than shuffled data (*p* < 0.001 for all positions, t-test with significant *p*-values for at least 3 consecutive positions). **d**, Angles between the decoding axes across maze positions for each variable, averaged across mice. Note that angles change fastest for task decoders (see Fig. 2f).

**Fig. S5.**
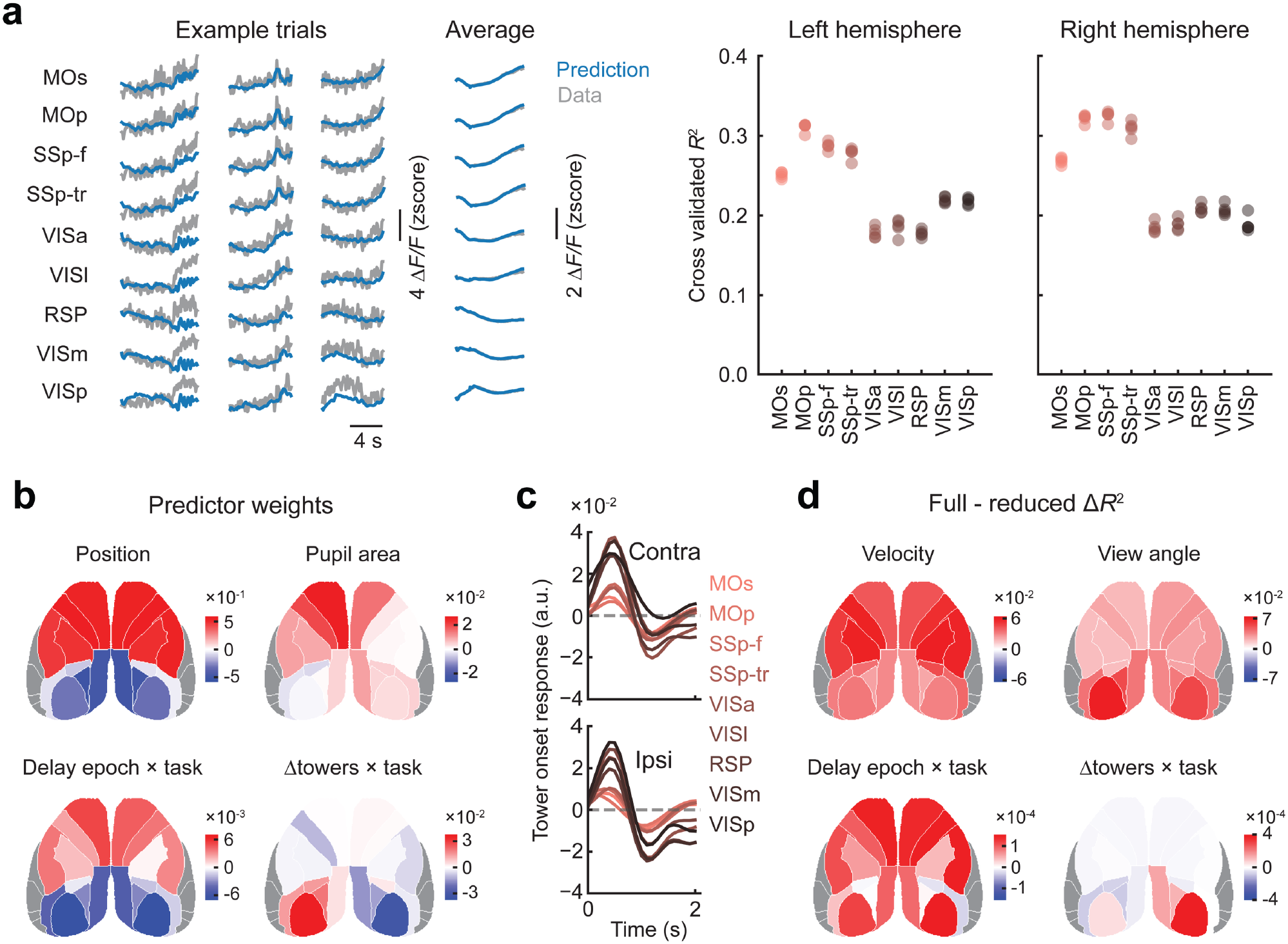
Further quantification of the linear encoding model of cholinergic activity. **a**, Left: Examples of single-trial and trial-averaged traces in the held-out test set, with model predictions. Right: Cross-validated model *R*^*2*^ per cortical region. Each data point corresponds to one cross-validation fold. **b**, Example maps of model coefficients by cortical area. **c**, Response kernels to contra- and ipsi-lateral towers for cholinergic axons in different cortical regions, estimated with the linear encoding model. Error estimates are omitted for clarity. **d**, Additional example maps of contributions of different classes of model predictors to explaining cholinergic activity, estimated as the change in the amount of cross-validated variance explained between the full model and a reduced model lacking that set of predictors and refitted to the data. See Table S1 for full quantification.

**Fig. S6.**
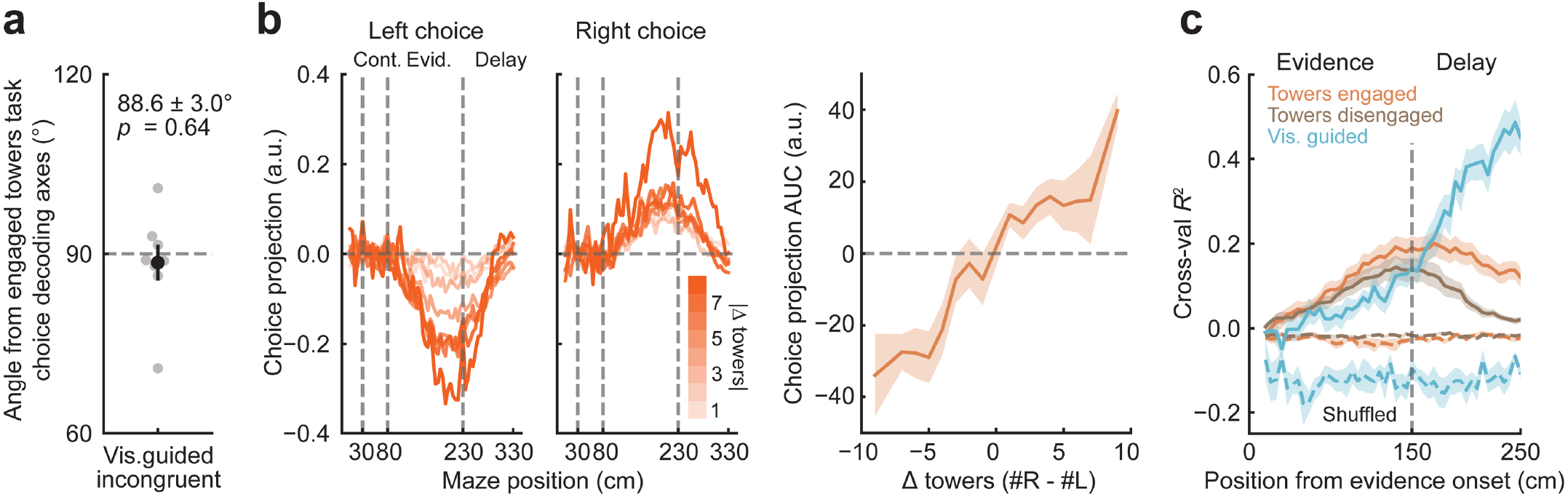
Additional quantification of evidence and choice decoding from cholinergic activity. **a**, Average angles between decoding axes for towers-task (engaged) and incongruent visually guided trials, computed separately for each animal within the evidence region. Gray data points: individual mice; black point and error bars: mean ± s.e.m. across mice (n = 8). Printed *p*-value is from a one-sided t-test. **b**, Analysis as in Fig. 3c, d, but performed on cholinergic activity after regressing out view angle (see Methods for details). Left: projections of cholinergic activity onto the choice-decoding axes for the towers task, separately for trials with different amounts of sensory evidence (|Δ towers|, color bar), averaged across mice. Right: AUC of the choice projections on the left, plotted as a function of sensory evidence for engaged trials in the towers task. Error shades: ± s.e.m. across mice (n = 8). **c**, Accuracy of linear-regression decoders trained to classify cumulative evidence from area-averaged cholinergic-axon activity in different parts of the maze, for engaged- and disengaged-strategy trials in the towers task, and incongruent visually guided trials. Error shades: ± s.e.m. across mice (n = 8).

**Fig. S7.**
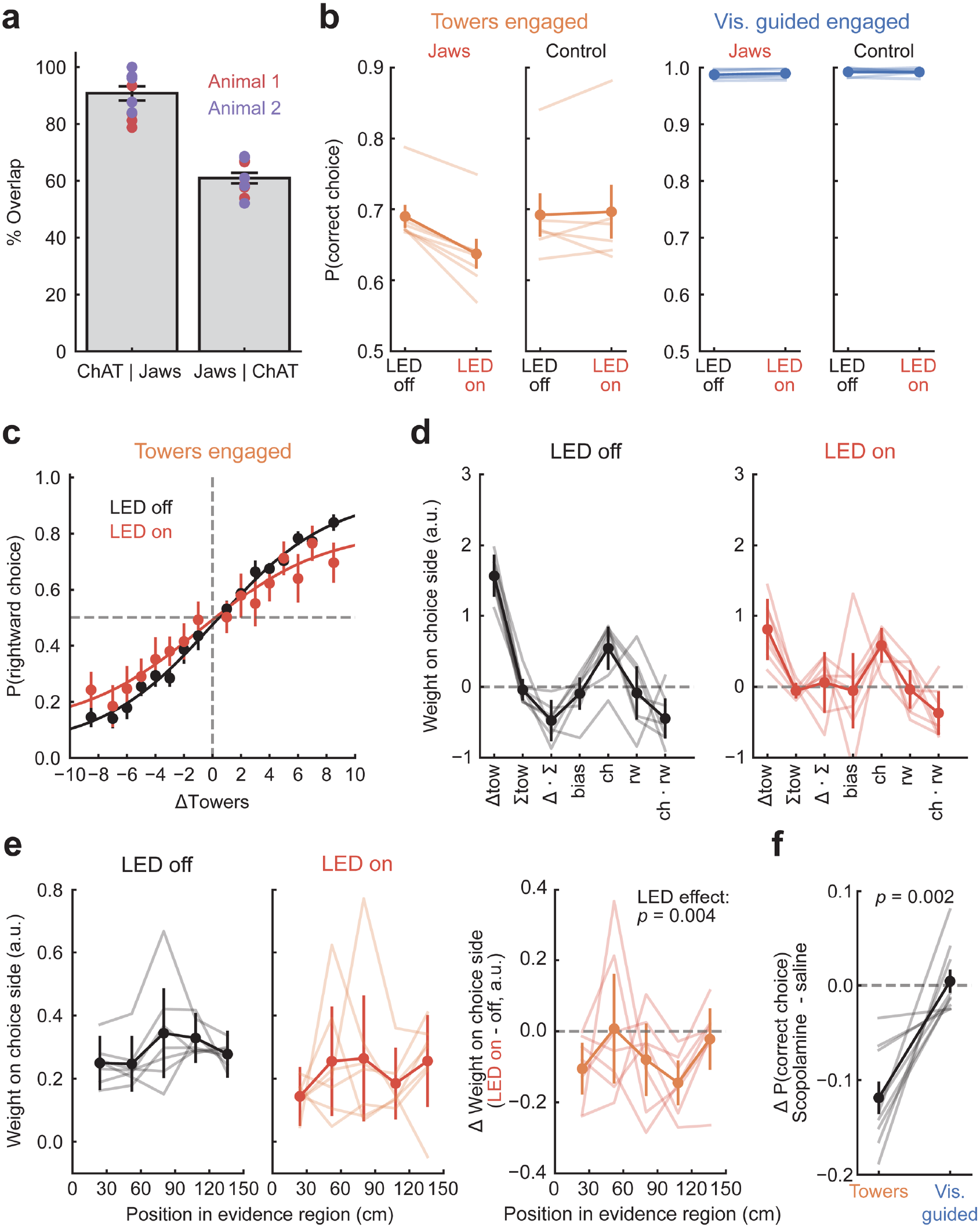
Additional quantification of the behavioral effects of perturbing cholinergic activity. **a**, Quantification of the specificity (ChAT | Jaws) and penetrance (Jaws | ChAT) of viral Jaws expression in basal forebrain cholinergic neurons. Data points correspond to individual fields of view, colored according to the animal they came from. Error bars: ± s.e.m. across fields of view. **b**, Overall performance of each task with and without optogenetic silencing, in Jaws-expressing (n = 7) and control (n = 6) mice. Thin lines: individual mice. Thick lines, circles and error bars: mean ± s.e.m. across mice. Refer to Fig. 4 for statistics. **c**, Psychometric curves computed separately for ‘LED on’ and ‘LED off’ trials during engaged strategies in the towers task, shown only for Jaws-expressing animals. Data points and error bars: mean ± s.e.m. across mice (n = 7). Curves: best-fitting sigmoids. **d**, Logistic-regression models as in Fig. 1e, fitted separately to ‘LED on’ and ‘LED off’ trials during engaged strategies in the towers task. Thin lines: random-effect estimates for individual mice, thick lines: population fixed-effect estimates, error bars: mean ± c.i.. Refer to Fig. 4 for statistics. **e**, Left and middle: coefficients from the logistic-regression model as in Fig. 1d, which estimates how mice weigh evidence from the evidence region during engaged-strategy towers-task trials, fitted separately to ‘LED on’ and ‘LED off’ trials. Conventions as in d. Right: subtraction of the coefficients on the left. Thin lines: individual mice (n = 7). Thick lines, circles and data points: mean ± s.e.m. across mice. Print *p*-value is from two-way mixed ANOVA with position being the within factor and LED effect being the between factor. **f**, Change in overall performance of each task between sessions in which mice received saline or the pan-muscarinic antagonist scopolamine. Thin lines: individual mice (n = 9). Thick line, circles, error bars: mean ± s.e.m. across mice. Print *p*-value is from paired t-test.

**Fig. S8.**
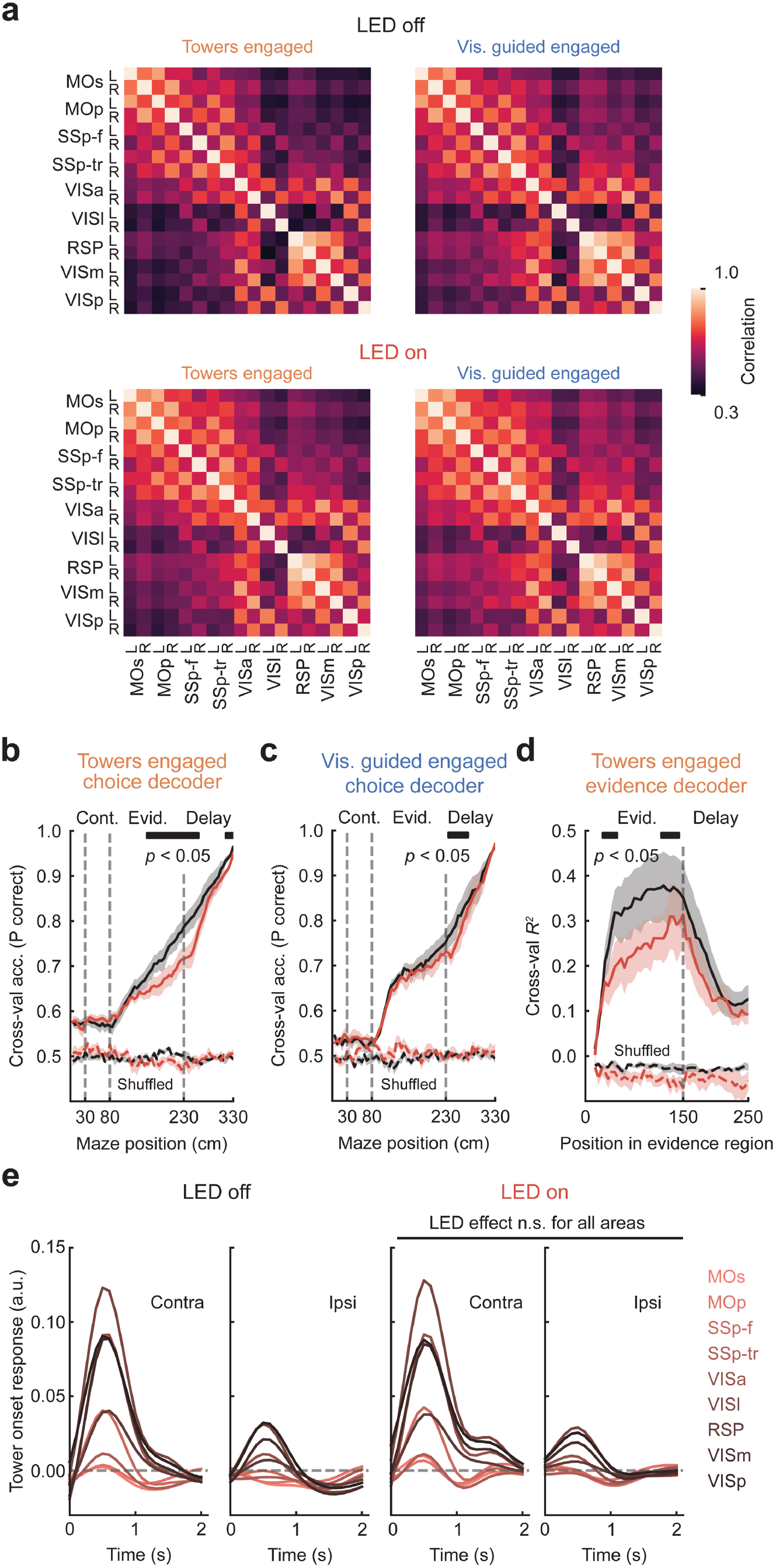
Additional quantification of the neural effects of perturbing cholinergic activity. **a**, Average pairwise correlations between excitatory activity of different cortical regions, shown separately for each task, and for control and cholinergic silencing trials. See Fig. 4 for statistics. **b**, Accuracy of choice decoders of excitatory activity during engaged strategies in the towers task, averaged within cortical regions, trained separately for each maze position and ‘LED on’ and ‘LED off’ trials. Error shades: ± s.e.m. across mice (n = 6). Bars on top indicate maze positions with a significant LED effect (*p* < 0.05 for at least 3 consecutive positions, t-test). **c**, Same as b, for engaged visually guided trials. **d**, Accuracy of linear-regression decoders of cumulative sensory evidence (Δ towers) during engaged towers-task trials. Conventions as in b. **e**, Response kernels to contra- and ipsi-lateral towers for excitatory activity in different cortical regions, estimated with the same linear encoding model from Fig. 2g on engaged towers-task trials with or without the LED × tower onset interaction. Error estimates are omitted for clarity. Removing the interaction term resulted in no significant reduction in model *R*^2^ with FDR correction.

**Table S1.**
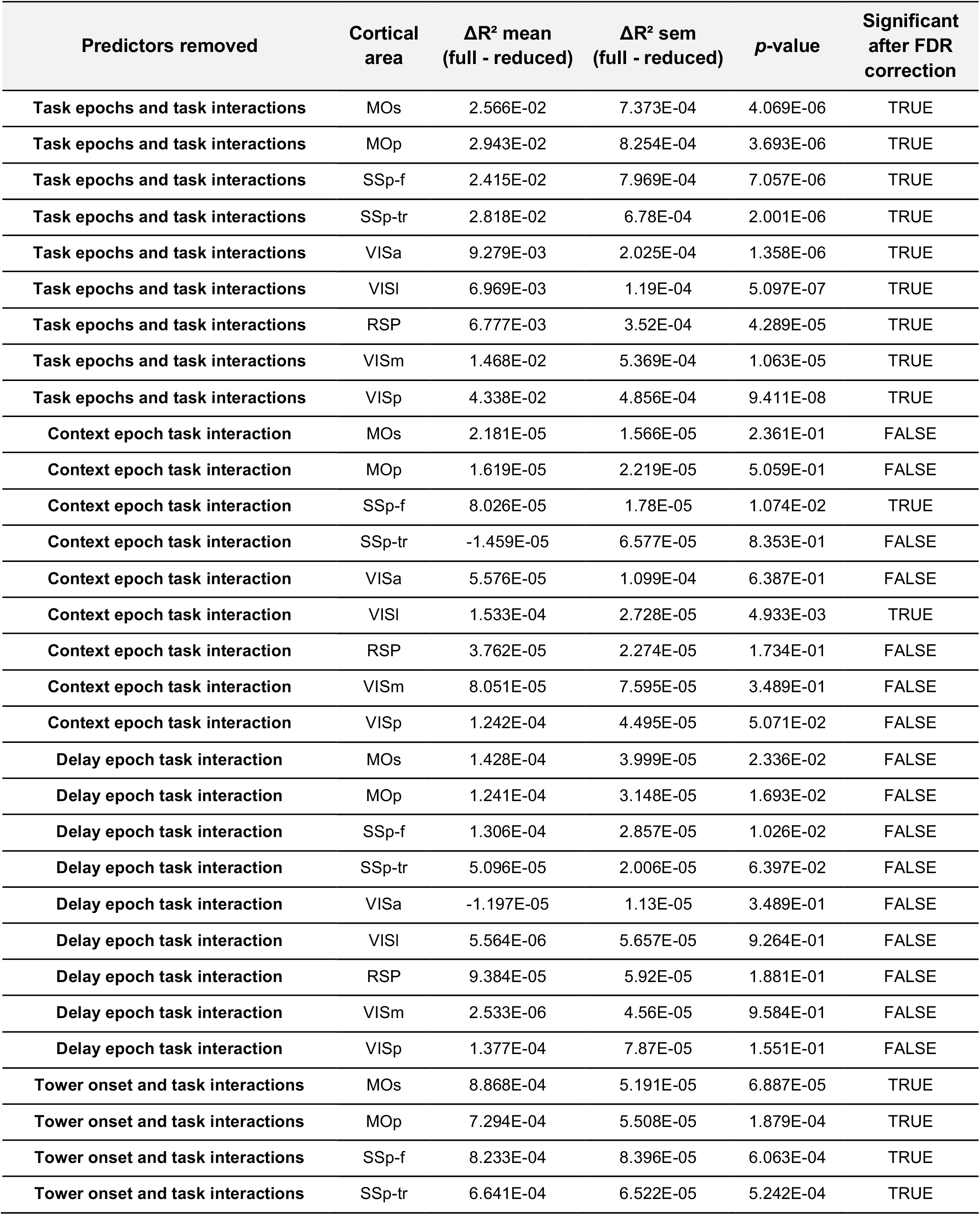
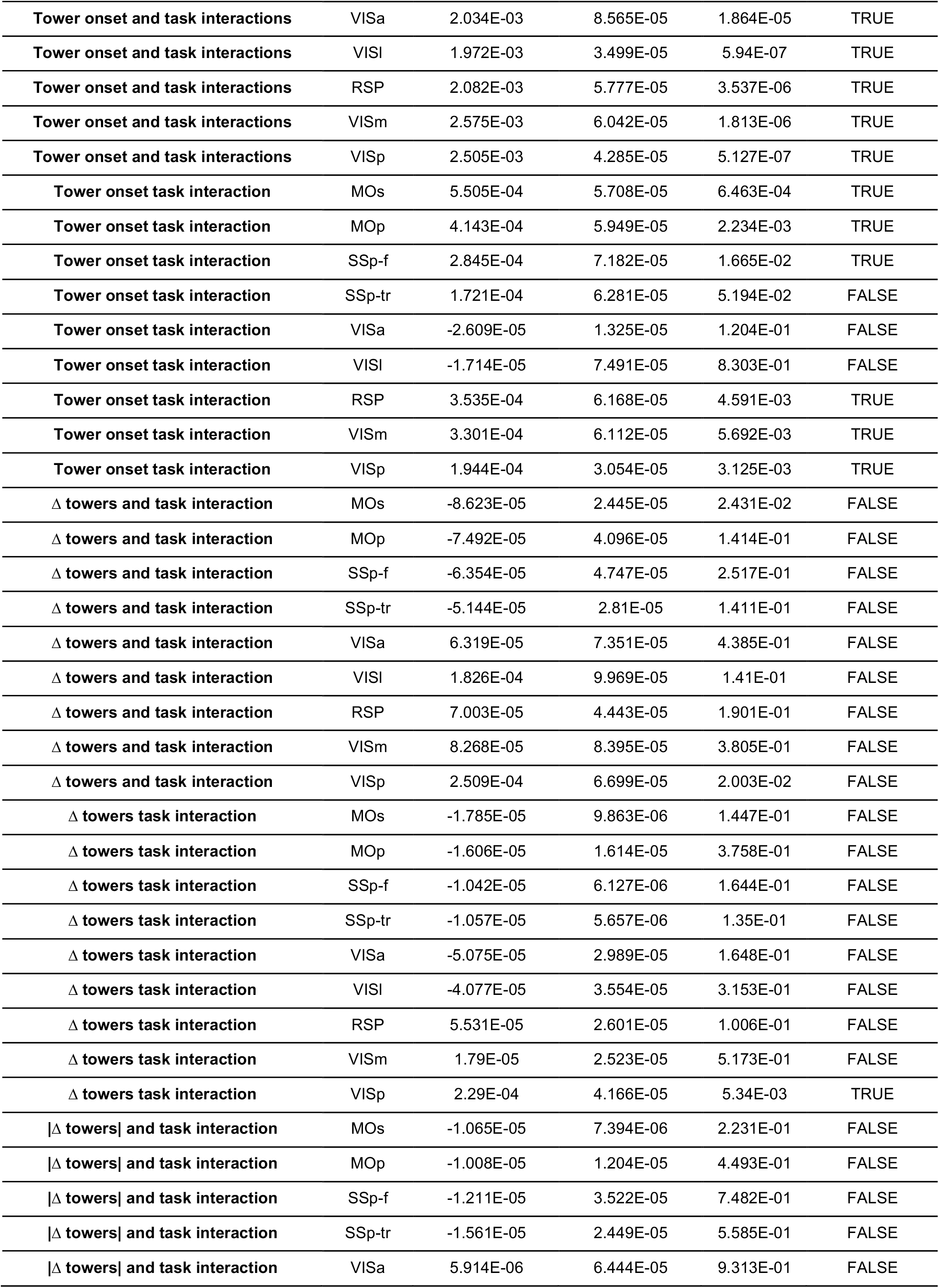
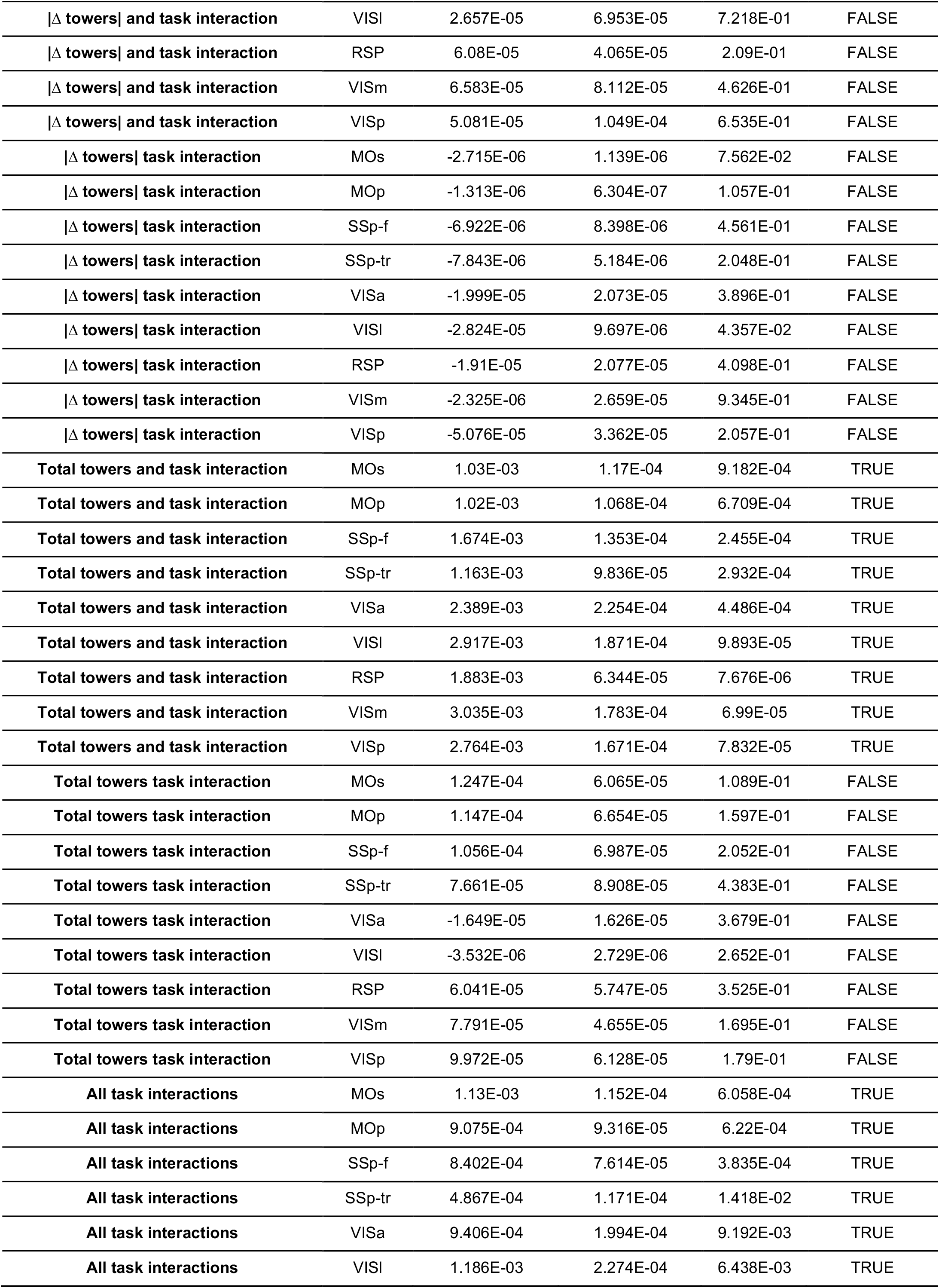
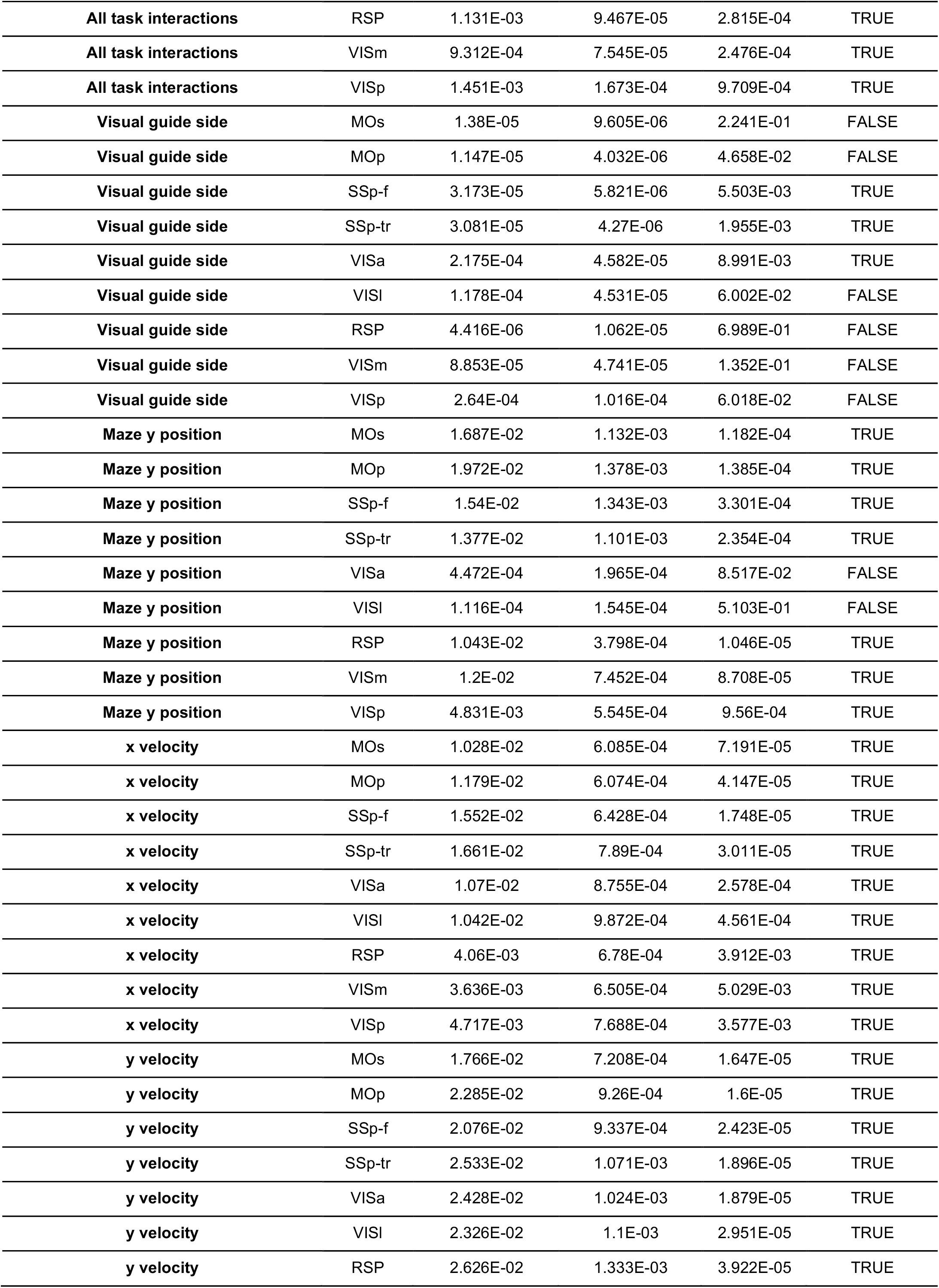
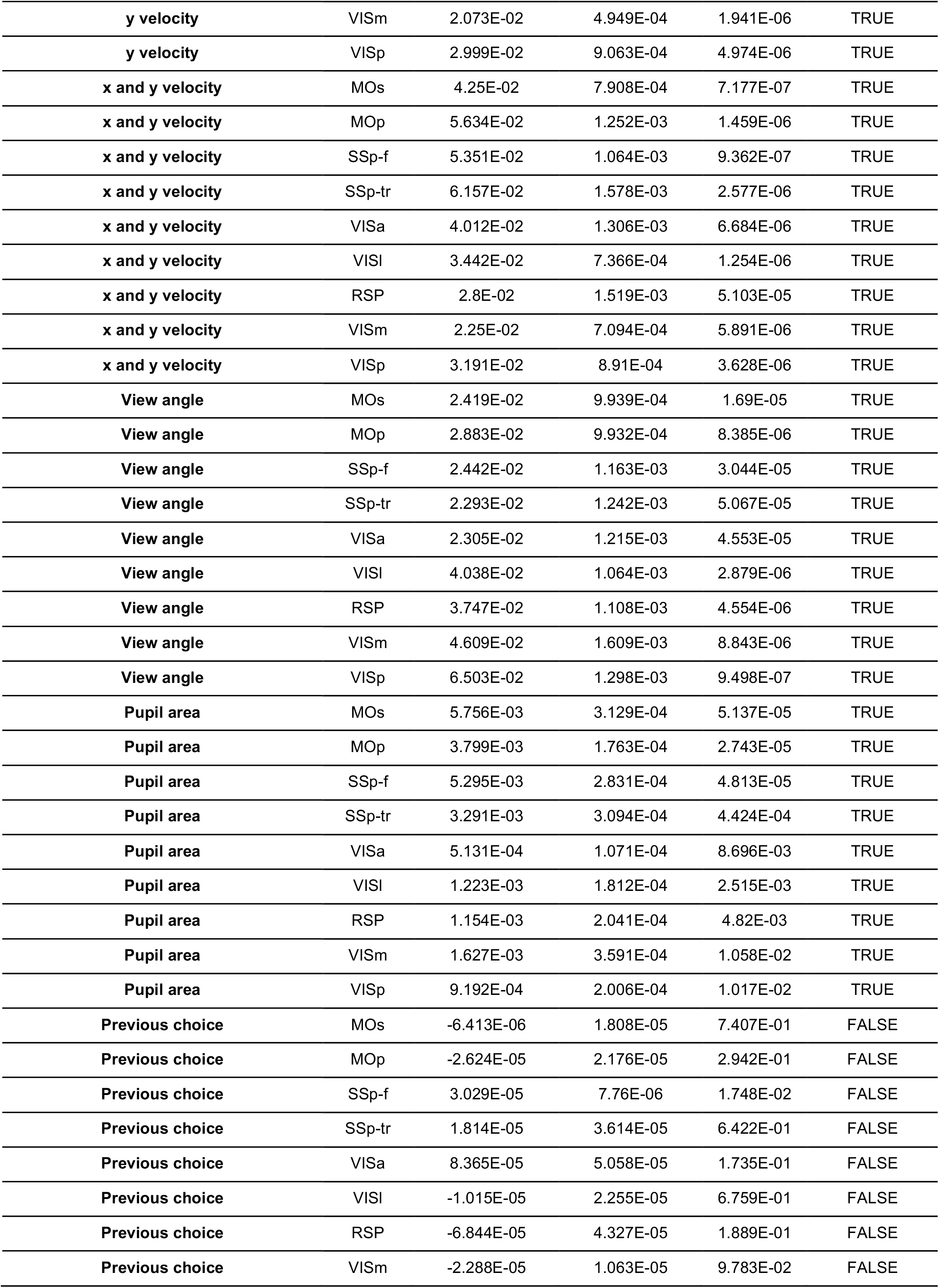
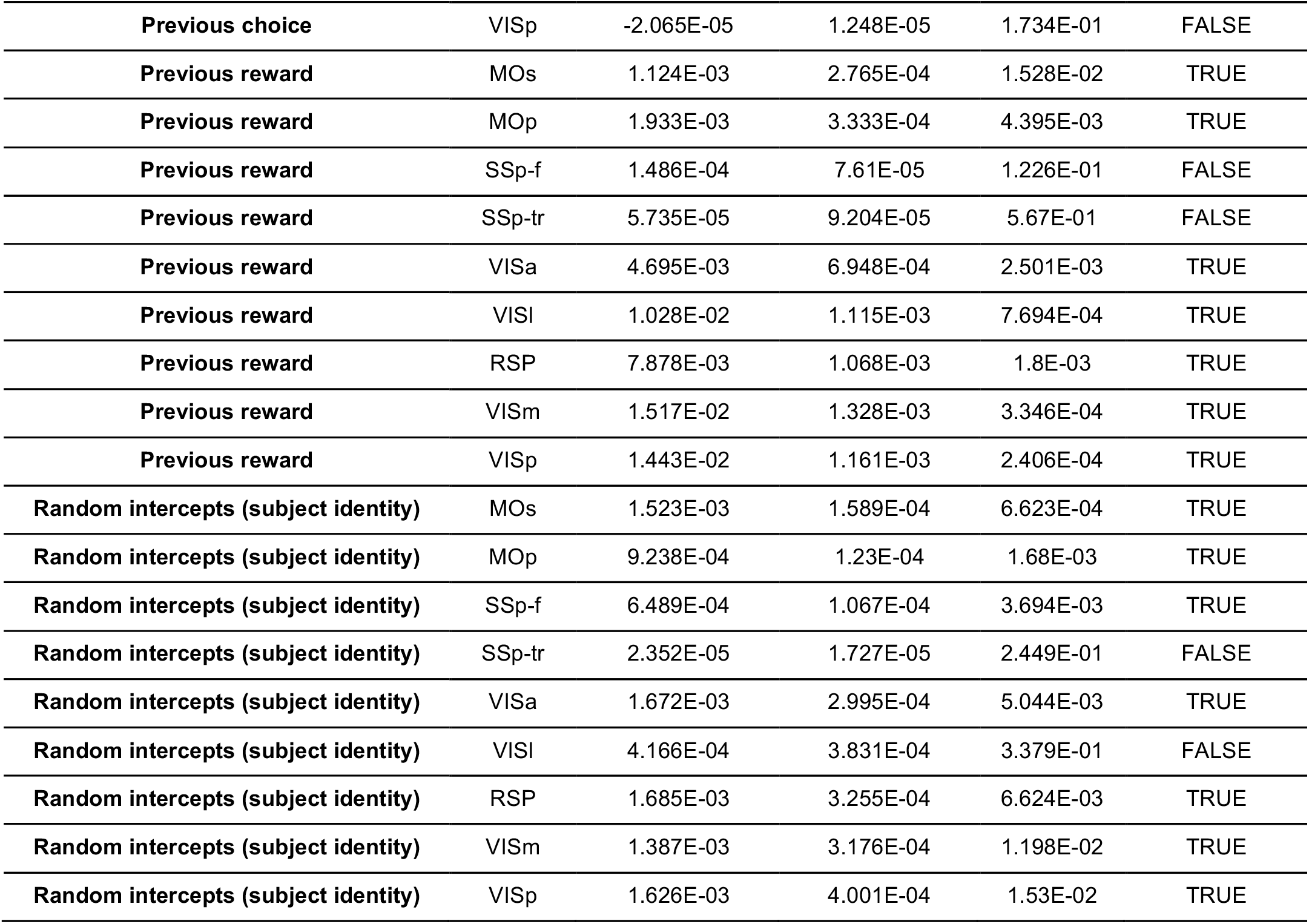
Statistics for predictor removals from the linear encoding model of cholinergic activity. For this analysis, the reduction in R^2^ was combined across hemispheres, and statistics were computed for the combined data.

**Table S2.**
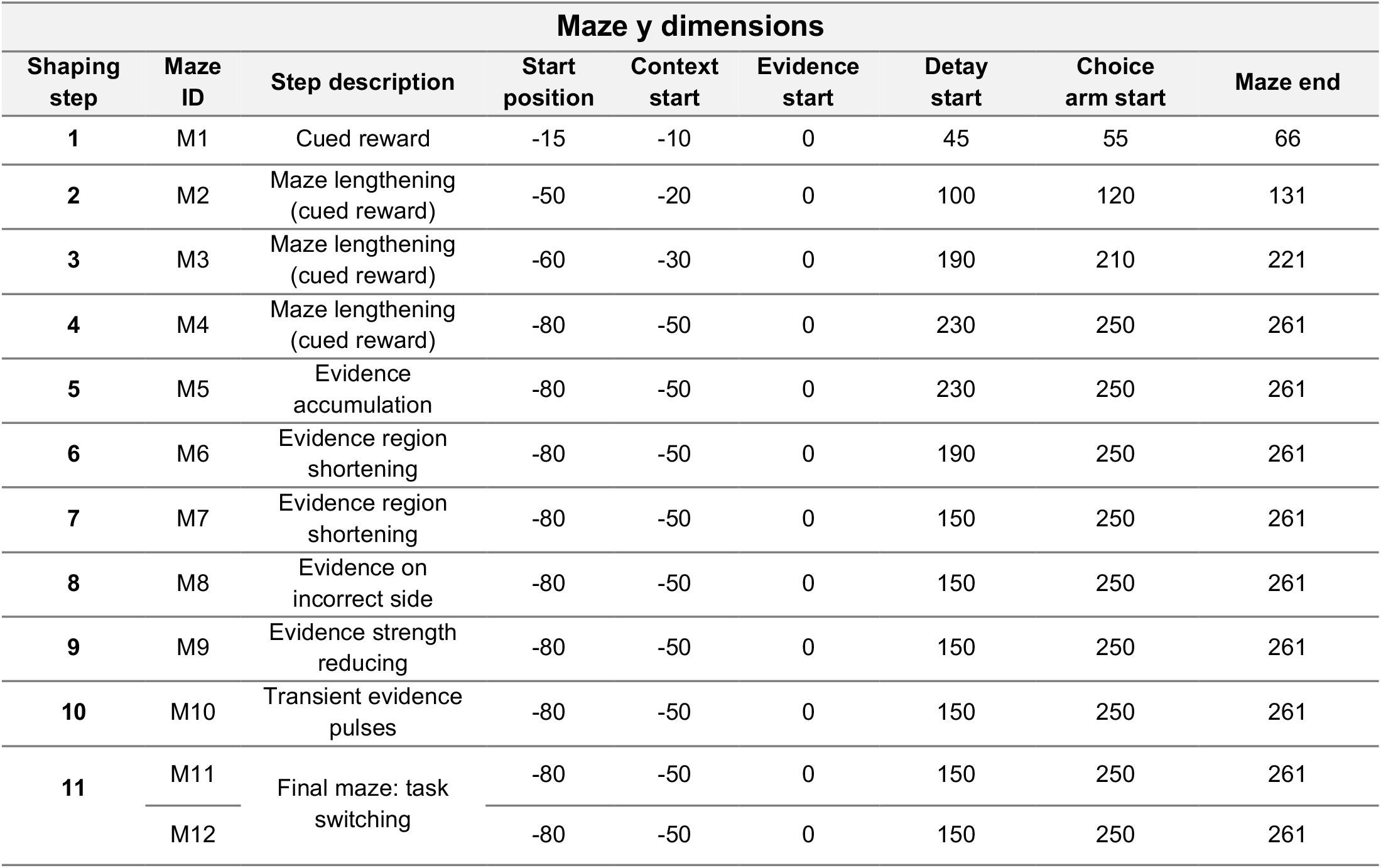
Maze longitudinal dimensions for each shaping step.

| Maze y dimensions |  |  |  |  |  |  |  |  |
| --- | --- | --- | --- | --- | --- | --- | --- | --- |
| Shaping step | Maze ID | Step description | Start position | Context start | Evidence start | Delay start | Choice arm start | Maze end |
| 1 | M1 | Cued reward | -15 | -10 | 0 | 45 | 55 | 66 |
| 2 | M2 | Maze lengthening (cued reward) | -50 | -20 | 0 | 100 | 120 | 131 |
| 3 | M3 | Maze lengthening (cued reward) | -60 | -30 | 0 | 190 | 210 | 221 |
| 4 | M4 | Maze lengthening (cued reward) | -80 | -50 | 0 | 230 | 250 | 261 |
| 5 | M5 | Evidence accumulation | -80 | -50 | 0 | 230 | 250 | 261 |
| 6 | M6 | Evidence region shortening | -80 | -50 | 0 | 190 | 250 | 261 |
| 7 | M7 | Evidence region shortening | -80 | -50 | 0 | 150 | 250 | 261 |
| 8 | M8 | Evidence on incorrect side | -80 | -50 | 0 | 150 | 250 | 261 |
| 9 | M9 | Evidence strength reducing | -80 | -50 | 0 | 150 | 250 | 261 |
| 10 | M10 | Transient evidence pulses | -80 | -50 | 0 | 150 | 250 | 261 |
| 11 | M11 | Final maze: task switching | -80 | -50 | 0 | 150 | 250 | 261 |
|  | M12 |  | -80 | -50 | 0 | 150 | 250 | 261 |

**Table S3.** Key task parameters for each shaping step. Maze switch indicates mazes into which mice may switch within a session. Before shaping step 11, this is triggered by a drop below the corresponding P(correct) threshold, computed over a 40-trial running window. In step 11, task switches become stochastic as described in Methods. Log(evidence ratio) is the natural logarithm of the ratio between Poisson rates for towers on rewarded and non-rewarded sides. Inf corresponds to towers only on the rewarded side. Inf pulse duration indicates permanently visible towers.

| Task parameters |  |  |  |  |  |  |  |
| --- | --- | --- | --- | --- | --- | --- | --- |
| Shaping step | Maze ID | Step description | Maze switch | P(correct) threshold for switch | log(evidence ratio) | Evidence pulse duration (ms) | Visual guide presence |
| 1 | M1 | Cued reward | N/A | N/A | inf | inf | True |
| 2 | M2 | Maze lengthening (cued reward) | N/A | N/A | inf | inf | True |
| 3 | M3 | Maze lengthening (cued reward) | N/A | N/A | inf | inf | True |
| 4 | M4 | Maze lengthening (cued reward) | N/A | N/A | inf | inf | True |
| 5 | M5 | Evidence accumulation | M4 | 0.70 | inf | inf | False |
| 6 | M6 | Evidence region shortening | M4 | 0.70 | inf | inf | False |
| 7 | M7 | Evidence region shortening | M4 | 0.70 | inf | inf | False |
| 8 | M8 | Evidence on incorrect side | M4 | 0.65 | 2.5 | inf | False |
| 9 | M9 | Evidence strength reducing | M7 | 0.60 | 1.6 | inf | False |
| 10 | M10 | Transient evidence pulses | M7 | 0.55 | 1.6 | 200 | False |
| 11 | M11 | Final maze: task switching | M12 | N/A | 1.2 | 200 | False |
|  | M12 |  | M11 | N/A | 1.2 | 200 | True |

**Table S4.**
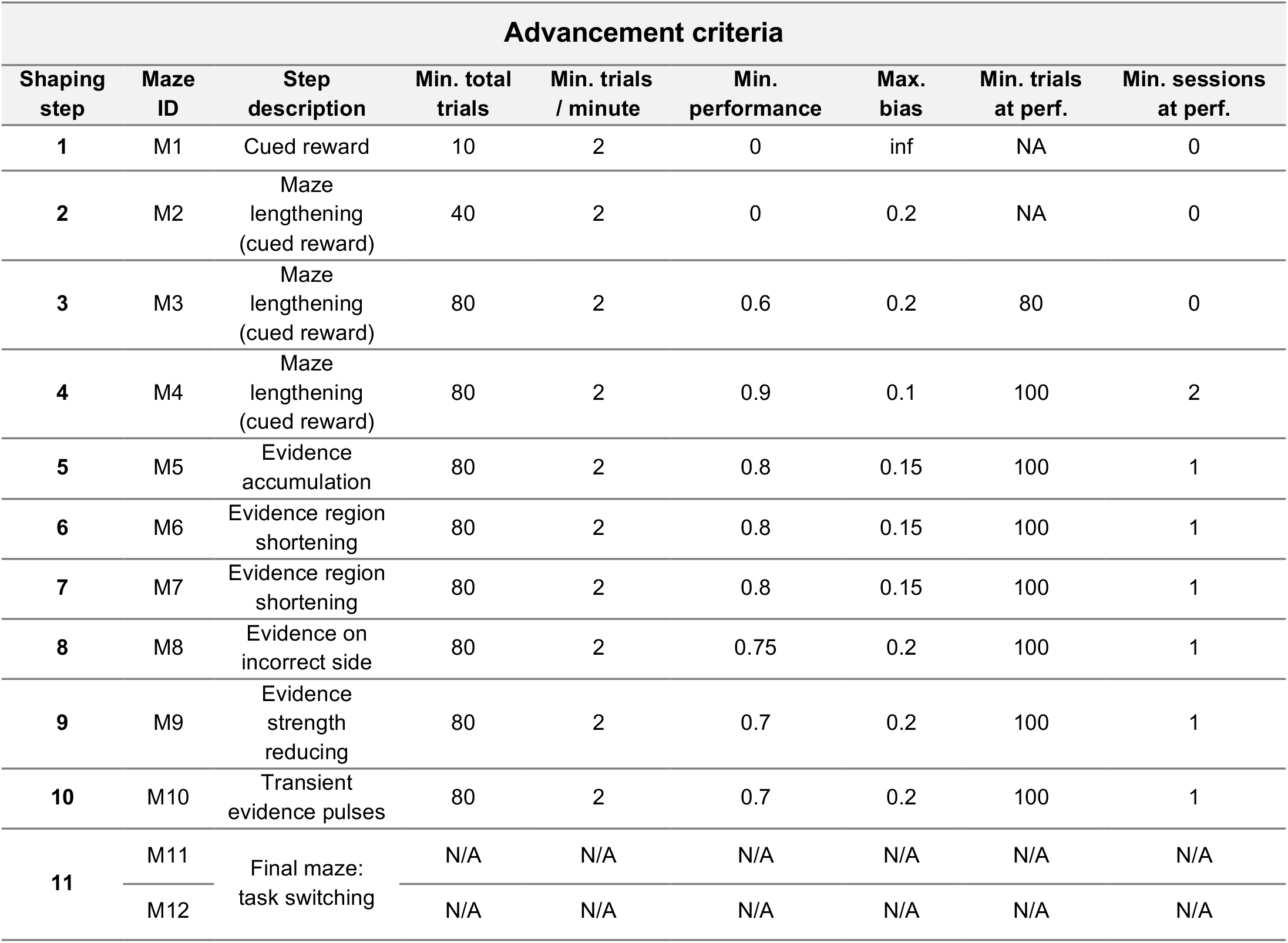
Advancement criteria for each shaping step. Shown are the maximum side bias and minimum performance required to advance to the following shaping step, and the minimum number of trials and sessions at performance criterion.

| Advancement criteria |  |  |  |  |  |  |  |  |
| --- | --- | --- | --- | --- | --- | --- | --- | --- |
| Shaping step | Maze ID | Step description | Min. total trials | Min. trials / minute | Min. performance | Max. bias | Min. trials at perf. | Min. sessions at perf. |
| 1 | M1 | Cued reward | 10 | 2 | 0 | inf | NA | 0 |
| 2 | M2 | Maze lengthening (cued reward) | 40 | 2 | 0 | 0.2 | NA | 0 |
| 3 | M3 | Maze lengthening (cued reward) | 80 | 2 | 0.6 | 0.2 | 80 | 0 |
| 4 | M4 | Maze lengthening (cued reward) | 80 | 2 | 0.9 | 0.1 | 100 | 2 |
| 5 | M5 | Evidence accumulation | 80 | 2 | 0.8 | 0.15 | 100 | 1 |
| 6 | M6 | Evidence region shortening | 80 | 2 | 0.8 | 0.15 | 100 | 1 |
| 7 | M7 | Evidence region shortening | 80 | 2 | 0.8 | 0.15 | 100 | 1 |
| 8 | M8 | Evidence on incorrect side | 80 | 2 | 0.75 | 0.2 | 100 | 1 |
| 9 | M9 | Evidence strength reducing | 80 | 2 | 0.7 | 0.2 | 100 | 1 |
| 10 | M10 | Transient evidence pulses | 80 | 2 | 0.7 | 0.2 | 100 | 1 |
| 11 | M11 | Final maze: task switching | N/A | N/A | N/A | N/A | N/A | N/A |
|  | M12 |  | N/A | N/A | N/A | N/A | N/A | N/A |

## References

1. Hernández, A. et al. Decoding a perceptual decision process across cortex. Neuron 66, 300–314 (2010).

2. Siegel, M., Buschman, T. J. & Miller, E. K. Cortical information flow during flexible sensorimotor decisions. Science 348, 1352–1355 (2015).

3. Allen, W. E. et al. Global representations of goal-directed behavior in distinct cell types of mouse neocortex. Neuron 94, 891–907.e6 (2017).

4. Dotson, N. M., Hoffman, S. J., Goodell, B. & Gray, C. M. Feature-based visual short-term memory is widely distributed and hierarchically organized. Neuron 99, 215– 226.e4 (2018).

5. Pinto, L. et al. Task-dependent changes in the large-scale dynamics and necessity of cortical regions. Neuron 104, 810–824.e9 (2019).

6. Steinmetz, N. A., Zatka-Haas, P., Carandini, M. & Harris, K. D. Distributed coding of choice, action and engagement across the mouse brain. Nature 576, 266–273 (2019).

7. Chen, S. et al. Brain-wide neural activity underlying memory-guided movement. Cell 187, 676–691.e16 (2024).

8. DePasquale, B., Brody, C. D. & Pillow, J. W. Neural population dynamics underlying evidence accumulation in multiple rat brain regions. Elife 13, e84955 (2024).

9. Khilkevich, A. et al. Brain-wide dynamics linking sensation to action during decision-making. Nature 634, 890–900 (2024).

10. Gilad, A., Gallero-Salas, Y., Groos, D. & Helmchen, F. Behavioral Strategy Determines Frontal or Posterior Location of Short-Term Memory in Neocortex. Neuron 99, 814–828.e7 (2018).

11. International Brain Laboratory et al. A brain-wide map of neural activity during complex behaviour. Nature 645, 177–191 (2025).

12. Yin, C. et al. Spontaneous movements and their relationship to neural activity fluctuate with latent engagement states. Neuron 113, 3048–3063.e5 (2025).

13. Musall, S. et al. Pyramidal cell types drive functionally distinct cortical activity patterns during decision-making. Nat. Neurosci. 26, 495–505 (2023).

14. Arlt, C. et al. Cognitive experience alters cortical involvement in goal-directed navigation. Elife 11, e76051 (2022).

15. Wilming, N., Murphy, P. R., Meyniel, F. & Donner, T. H. Large-scale dynamics of perceptual decision information across human cortex. Nat. Commun. 11, 5109 (2020).

16. Ito, T., Hearne, L. J. & Cole, M. W. A cortical hierarchy of localized and distributed processes revealed via dissociation of task activations, connectivity changes, and intrinsic timescales. Neuroimage 221, 117141 (2020).

17. Krueger, P. M. et al. Evidence accumulation detected in BOLD signal using slow perceptual decision making. J. Neurosci. Methods 281, 21–32 (2017).

18. Zaborszky, L., van den Pol, A. & Gyengési, E. The Basal Forebrain Cholinergic Projection System in Mice. In The mouse nervous system (eds. Watson, C., Paxinos, G. & Puelles, L.) 684–709 (Elsevier, San Diego, 2011).

19. Muñoz, W. & Rudy, B. Spatiotemporal specificity in cholinergic control of neocortical function. Curr. Opin. Neurobiol. 26, 149–160 (2014).

20. Záborszky, L. et al. Specific basal forebrain-cortical cholinergic circuits coordinate cognitive operations. J. Neurosci. 38, 9446–9458 (2018).

21. Laplante, F., Morin, Y., Quirion, R. & Vaucher, E. Acetylcholine release is elicited in the visual cortex, but not in the prefrontal cortex, by patterned visual stimulation: a dual in vivo microdialysis study with functional correlates in the rat brain. Neuroscience 132, 501–510 (2005).

22. Parikh, V., Kozak, R., Martinez, V. & Sarter, M. Prefrontal acetylcholine release controls cue detection on multiple timescales. Neuron 56, 141–154 (2007).

23. Pinto, L. et al. Fast modulation of visual perception by basal forebrain cholinergic neurons. Nat. Neurosci. 16, 1857–1863 (2013).

24. Robert, B. et al. A functional topography within the cholinergic basal forebrain for encoding sensory cues and behavioral reinforcement outcomes. Elife 10, e69514 (2021).

25. Lohani, S. et al. Spatiotemporally heterogeneous coordination of cholinergic and neocortical activity. Nat. Neurosci. 25, 1706–1713 (2022).

26. Yu, B. et al. Cholinergic feedback for modality- and context-specific modulation of sensory representations. Science 388, 1324–1329 (2025).

27. Buzsaki, G. et al. Nucleus basalis and thalamic control of neocortical activity in the freely moving rat. J. Neurosci. 8, 4007–4026 (1988).

28. Reimer, J. et al. Pupil fluctuations track rapid changes in adrenergic and cholinergic activity in cortex. Nat. Commun. 7, 13289 (2016).

29. Xu, M. et al. Basal forebrain circuit for sleep-wake control. Nat. Neurosci. 18, 1641– 1647 (2015).

30. Collins, L., Francis, J., Emanuel, B. & McCormick, D. A. Cholinergic and noradrenergic axonal activity contains a behavioral-state signal that is coordinated across the dorsal cortex. Elife 12, e81826 (2023).

31. Zhu, F., Elnozahy, S., Lawlor, J. & Kuchibhotla, K. V. The cholinergic basal forebrain provides a parallel channel for state-dependent sensory signaling to auditory cortex. Nat. Neurosci. 26, 810–819 (2023).

32. Li, B. et al. Circuit mechanism for suppression of frontal cortical ignition during NREM sleep. Cell 186, 5739–5750.e17 (2023).

33. Yogesh, B. & Keller, G. B. Cholinergic input to mouse visual cortex signals a movement state and acutely enhances layer 5 responsiveness. Elife 12, e89986.5 (2024).

34. Kozak, R., Bruno, J. P. & Sarter, M. Augmented prefrontal acetylcholine release during challenged attentional performance. Cereb. Cortex 16, 9–17 (2006).

35. Herrero, J. L. et al. Acetylcholine contributes through muscarinic receptors to attentional modulation in V1. Nature 454, 1110–1114 (2008).

36. Goard, M. & Dan, Y. Basal forebrain activation enhances cortical coding of natural scenes. Nat. Neurosci. 12, 1444–1449 (2009).

37. Eggermann, E., Kremer, Y., Crochet, S. & Petersen, C. C. H. Cholinergic signals in mouse barrel cortex during active whisker sensing. Cell Rep. 9, 1654–1660 (2014).

38. Harrison, T. C., Pinto, L., Brock, J. R. & Dan, Y. Calcium imaging of basal forebrain activity during innate and learned behaviors. Front. Neural Circuits 10, 36 (2016).

39. Laszlovszky, T. et al. Distinct synchronization, cortical coupling and behavioral function of two basal forebrain cholinergic neuron types. Nat. Neurosci. 23, 992– 1003 (2020).

40. Hasselmo, M. E. & Giocomo, L. M. Cholinergic modulation of cortical function. J. Mol. Neurosci. 30, 133–135 (2006).

41. Kuchibhotla, K. V. et al. Parallel processing by cortical inhibition enables context-dependent behavior. Nat. Neurosci. 1–14 (2016).

42. Bauer, M. et al. Cholinergic enhancement of visual attention and neural oscillations in the human brain. Curr. Biol. 22, 397–402 (2012).

43. Rokem, A., Landau, A. N., Garg, D., Prinzmetal, W. & Silver, M. A. Cholinergic enhancement increases the effects of voluntary attention but does not affect involuntary attention. Neuropsychopharmacology 35, 2538–2544 (2010).

44. Lee, A. M. et al. Identification of a brainstem circuit regulating visual cortical state in parallel with locomotion. Neuron 83, 455–466 (2014).

45. Fu, Y. et al. A Cortical Circuit for Gain Control by Behavioral State. Cell 156, 1139– 1152 (2014).

46. Nelson, A. & Mooney, R. The Basal Forebrain and Motor Cortex Provide Convergent yet Distinct Movement-Related Inputs to the Auditory Cortex. Neuron 1–15 (2016).

47. Khalighinejad, N. et al. A basal forebrain-cingulate circuit in macaques decides it is time to act. Neuron 105, 370–384.e8 (2020).

48. Chubykin, A. A., Roach, E. B., Bear, M. F. & Shuler, M. G. H. A cholinergic mechanism for reward timing within primary visual cortex. Neuron 77, 723–735 (2013).

49. Hangya, B., Ranade, S. P., Lorenc, M. & Kepecs, A. Central cholinergic neurons are rapidly recruited by reinforcement feedback. Cell 162, 1155–1168 (2015).

50. Froemke, R. C., Merzenich, M. M. & Schreiner, C. E. A synaptic memory trace for cortical receptive field plasticity. Nature 450, 425–429 (2007).

51. Kilgard, M. P. & Merzenich, M. M. Cortical map reorganization enabled by nucleus basalis activity. Science 279, 1714–1718 (1998).

52. Wang, H. et al. Frontal noradrenergic and cholinergic transients exhibit distinct spatiotemporal dynamics during competitive decision-making. Sci. Adv. 11, eadr9916 (2025).

53. Wu, H., Williams, J. & Nathans, J. Complete morphologies of basal forebrain cholinergic neurons in the mouse. Elife 3, e02444 (2014).

54. Gielow, M. R. & Zaborszky, L. The input-output relationship of the cholinergic basal forebrain. Cell Rep. 18, 1817–1830 (2017).

55. Do, J. P. et al. Cell type-specific long-range connections of basal forebrain circuit. Elife 5, e13214 (2016).

56. Bloem, B. et al. Topographic mapping between basal forebrain cholinergic neurons and the medial prefrontal cortex in mice. J. Neurosci. 34, 16234–16246 (2014).

57. Li, X. et al. Generation of a whole-brain atlas for the cholinergic system and mesoscopic projectome analysis of basal forebrain cholinergic neurons. Proc. Natl. Acad. Sci. U. S. A. 115, 415–420 (2018).

58. Huppé-Gourgues, F., Jegouic, K. & Vaucher, E. Topographic organization of cholinergic innervation from the basal forebrain to the visual cortex in the rat. Front. Neural Circuits 12, 19 (2018).

59. Chen, Z. et al. Whole-brain mapping of basal forebrain cholinergic neurons reveals a long-range reciprocal input-output loop between distinct subtypes. Sci. Adv. 11, eadt1617 (2025).

60. Knudstrup, S. G. et al. Visual stimulation drives retinotopic acetylcholine release in the mouse visual cortex. bioRxivorg (2024) doi:10.1101/2024.02.04.578821.

61. Pinto, L. et al. An accumulation-of-evidence task using visual pulses for mice navigating in virtual reality. Front. Behav. Neurosci. 12, 36 (2018).

62. Minces, V., Pinto, L., Dan, Y. & Chiba, A. A. Cholinergic shaping of neural correlations. Proc. Natl. Acad. Sci. U. S. A. 114, 5725–5730 (2017).

63. Turchi, J. et al. The Basal Forebrain Regulates Global Resting-State fMRI Fluctuations. Neuron 97, 940–952.e4 (2018).

64. Pfeffer, T. et al. Circuit mechanisms for the chemical modulation of cortex-wide network interactions and behavioral variability. Sci. Adv. 7, eabf5620 (2021).

65. Shine, J. M. Neuromodulatory influences on integration and segregation in the brain. Trends Cogn. Sci. 23, 572–583 (2019).

66. Calhoun, A. J., Pillow, J. W. & Murthy, M. Author Correction: Unsupervised identification of the internal states that shape natural behavior. Nat. Neurosci. 23, 293 (2020).

67. Ashwood, Z. C. et al. Mice alternate between discrete strategies during perceptual decision-making. Nat. Neurosci. 25, 201–212 (2022).

68. Bolkan, S. S. et al. Opponent control of behavior by dorsomedial striatal pathways depends on task demands and internal state. Nat. Neurosci. 25, 345–357 (2022).

69. Lee, S.-H. & Dan, Y. Neuromodulation of brain states. Neuron 76, 209–222 (2012).

70. Harris, K. D. & Thiele, A. Cortical state and attention. Nat. Rev. Neurosci. 12, 509– 523 (2011).

71. Monsell, S. Task switching. Trends Cogn. Sci. 7, 134–140 (2003).

72. Yu, A. J. & Dayan, P. Uncertainty, neuromodulation, and attention. Neuron 46, 681– 692 (2005).

73. Aston-Jones, G. & Cohen, J. D. An integrative theory of locus coeruleus-norepinephrine function: adaptive gain and optimal performance. Annu. Rev. Neurosci. 28, 403–450 (2005).

74. Kiani, R., Cueva, C. J., Reppas, J. B. & Newsome, W. T. Dynamics of Neural Population Responses in Prefrontal Cortex Indicate Changes of Mind on Single Trials. Curr. Biol. 24, 1542–1547 (2014).

75. van den Brink, R. L. et al. Flexible sensory-motor mapping rules manifest in correlated variability of stimulus and action codes across the brain. Neuron 111, 571–584.e9 (2023).

76. Bondy, A. G. et al. Coordinated cross-brain activity during accumulation of sensory evidence and decision commitment. bioRxiv (2024) doi:10.1101/2024.08.21.609044.

77. Costa, R. M. et al. Cognitive processes are disentangled at cortex-wide scales. bioRxiv 2025.07.24.666672 (2025) doi:10.1101/2025.07.24.666672.

78. Toso, A., Arazi, A., de la Rocha, J., Tsetsos, K. & Donner, T. H. Competing neural decision variables in human frontal cortex shape decision confidence. bioRxiv 2025.10.28.684827 (2025) doi:10.1101/2025.10.28.684827.

79. Zhang, Y. et al. Fast and sensitive GCaMP calcium indicators for imaging neural populations. Nature 615, 884–891 (2023).

80. Gold, J. I. & Shadlen, M. N. The neural basis of decision making. Annu. Rev. Neurosci. 30, 535–574 (2007).

81. Brody, C. D. & Hanks, T. D. Neural underpinnings of the evidence accumulator. Curr. Opin. Neurobiol. 37, 149–157 (2016).

82. Mante, V., Sussillo, D., Shenoy, K. V. & Newsome, W. T. Context-dependent computation by recurrent dynamics in prefrontal cortex. Nature 503, 78–84 (2013).

83. Pagan, M. et al. Individual variability of neural computations underlying flexible decisions. Nature 639, 421–429 (2025).

84. Diamanti, E. (mika) et al. Working memory expands shared task representations in cortex. bioRxiv (2025) doi:10.1101/2025.09.29.679345.

85. Golmayo, L., Nunez, A. & Zaborszky, L. Electrophysiological evidence for the existence of a posterior cortical-prefrontal-basal forebrain circuitry in modulating sensory responses in visual and somatosensory rat cortical areas. Neuroscience 119, 597–609 (2003).

86. Hasselmo, M. E. & McGaughy, J. High acetylcholine levels set circuit dynamics for attention and encoding and low acetylcholine levels set dynamics for consolidation. Prog. Brain Res. 145, 207–231 (2004).

87. Disney, A. A., Aoki, C. & Hawken, M. J. Gain modulation by nicotine in macaque v1. Neuron 56, 701–713 (2007).

88. Roberts, M. J. et al. Acetylcholine dynamically controls spatial integration in marmoset primary visual cortex. J. Neurophysiol. 93, 2062–2072 (2005).

89. Goldman, M. S. Memory without feedback in a neural network. Neuron 61, 621–634 (2009).

90. Scott, B. B. et al. Fronto-parietal Cortical Circuits Encode Accumulated Evidence with a Diversity of Timescales. Neuron 95, 385–398.e5 (2017).

91. Pinto, L., Tank, D. W. & Brody, C. D. Multiple timescales of sensory-evidence accumulation across the dorsal cortex. Elife 11, e70263 (2022).

92. Aronov, D. & Tank, D. W. Engagement of Neural Circuits Underlying 2D Spatial Navigation in a Rodent Virtual Reality System. Neuron 84, 442–456 (2014).

93. Nath, T. et al. Using DeepLabCut for 3D markerless pose estimation across species and behaviors. Nat. Protoc. 14, 2152–2176 (2019).

94. Biderman, D. et al. Lightning Pose: improved animal pose estimation via semi-supervised learning, Bayesian ensembling, and cloud-native open-source tools. bioRxivorg (2023) doi:10.1101/2023.04.28.538703.

95. Benjamini, Y. & Hochberg, Y. Controlling the false discovery rate: a practical and powerful approach to multiple testing. J. R. Stat. Soc. Series B Stat. Methodol. (1995).

96. Jean-Richard-Dit-Bressel, P., Clifford, C. W. G. & McNally, G. P. Analyzing event-related transients: Confidence intervals, permutation tests, and consecutive thresholds. Front. Mol. Neurosci. 13, 14 (2020).

